# HER2 Cancer Protrusion Growth Signaling Regulated by Unhindered, Localized Filopodial Dynamics

**DOI:** 10.1101/654988

**Authors:** Wai Yan Lam, Yi Wang, Barmak Mostofian, Danielle Jorgens, Sunjong Kwon, Koei Chin, M. Alexandra Carpenter, Thomas Jacob, Katie Heiser, Anurag Agrawal, Jing Wang, Xiaolin Nan, Young Hwan Chang, Daniel M. Zuckerman, Joe Gray, Marcel Bruchez, Keith A. Lidke, Tania Q. Vu

## Abstract

Protrusions are plasma membrane extensions that are found in almost every cell in the human body. Cancer cell filopodial and lamellipodial protrusions play key roles in the integral processes of cell motility and signaling underlying tumor invasion and metastasis. HER2 (ErbB-2) is overexpressed in diverse types of tumors and regulates PI3K-pathway-mediated protrusion growth. It is known that HER2 resides at breast cancer cell protrusions, but how protrusion-based HER2 spatiotemporal dynamics shape cancer signaling is unclear. Here, we study how HER2 location and motion regulate protrusion signaling and growth using quantitative spatio-temporal molecular imaging approaches. Our data highlight morphologically-segregated features of filopodial and lamellipodial protrusions, in *in vitro* 2D breast cancer cells and *in vivo* intact breast tumor. Functional-segregation parallels morphological-segregation, as HER2 and its activated downstream pAKT-PI3K signaling remain spatially-localized at protrusions, provoking new protrusion growth proximal to sites of HER2 activation. HER2 in SKBR3 breast cancer cell filopodia exhibits fast, linearly-directed motion that is distinct from lamellipodia and non-protrusion subcellular regions (∼3-4 times greater diffusion constant, rapid speeds of 2-3 um2/s). Surprisingly, filopodial HER2 motion is passive, requiring no active energy sources. Moreover, while HER2 motion in lamellipodia and non-protrusion regions show hindered diffusion typical of membrane proteins, HER2 diffuses freely within filopodia. We conclude that HER2 activation, propagation, and functional protrusion growth is a local process in which filopodia have evolved to exploit Brownian thermal fluctuations within a barrier-free nanostructure to transduce rapid signaling. These results support the importance of developing filopodia and other protrusion-targeted strategies for cancer.

## Introduction

Cell plasma membrane protrusions play important roles in transducing signaling and in promoting cell motility, processes that are essential to many fundamental physiological functions (e.g. morphogenesis, wound healing, and immune surveillance) [1-3]. Aberrant protrusion regulation is a basis for a diversity of pathologies affecting a wide range of physiological functions (e.g. pancreatic disease, retinal degeneration, cognitive defects, obesity) and tissue types (e.g. brain, kidney, and lung) [4-6]. In cancer, dysregulated protrusion function leads to an overabundance of filopodia and lamellipodia, two main types of actin-based protrusions that form at the leading edge of migrating tumor cells [3, 7, 8]. Notably, filopodial and lamellipodial overabundance is correlated with tumor cell invasiveness, the primary cause of death in cancer patients [3, 7, 9-11]. Understanding how filopodia and lammelipodia regulate protrusion growth signaling is key to understanding how to better control detrimental cancer signaling.

HER2 is a membrane receptor whose overexpression is typically of epithelial origin and present in a diversity of tumor tissues (e.g. breast, ovarian, prostate, pancreatic, gastric), contributing to excessive signaling that underlies tumor growth and invasion [12-17]). Because HER2 is an effective therapeutic target in breast and gastric cancer, and because patients of diverse cancers have responded to anti-HER2 therapies, there is clinical and fundamental importance in understanding HER2 regulation and trafficking [13, 17]. Signaling is a process that is dependent on coordination of location and timing of molecules. Intriguingly, HER2 is associated with membrane protrusions [18]. Past studies of HER2-and other ERB family receptor biology-have largely not considered spatial locale as an aspect governing in receptor regulation. Recent studies however have highlighted the importance of HER2 signaling in the formation and regulation of protrusions in breast cancer, as well as proposed new mechanisms of HER2 membrane regulation at protrusions [19-22]. The dynamics of EGFR (ERB-1), a receptor that is closely related to HER2 has been described and found, interestingly, to move in a retrograde fashion via active cytoskeletal transport along filopodia, and internalized for signaling at the cell body in live cells [23]. Furthermore, the *in vivo* HER2 trafficking to tumors [24]; as well, the dimerization dynamics of HER2 have been described [23, 25]. Strikingly, knowledge of the spatiotemporal dynamics of HER2 and its role in shaping protrusion growth signaling remains unclear.

Here we address the question of how the location and motion of HER2 may shape cancer cell protrusion signaling. We focus on elucidating the spatiotemporal dynamics HER2 in filopodial and lamellipodial protrusions, and non-protrusion regions of breast cancer cells using a combination of subcellular and molecular imaging approaches that we have developed to mitigate challenges in measuring the quantity and motion of signaling molecules in small protrusion structures. We study the morphology of breast cancer cell protrusions in 2D in vitro breast cancer cells and in 3D intact tumor, and quantitate localized concentrations of HER2 and pAKT complexes in protrusion and non-protrusion regions of 2D in vitro breast cancer cells using 3D FIB-SEM methodologies [26] and STORM/molecular counting techniques [27, 28]. Using new compact monovalent HER2 affibody-QD probes that we developed in this current study, along with our single particle analytical methodologies [29-34], we tracked and analyzed the particle dynamics of single HER2 complexes over extended durations (several minutes). These long-duration trajectories enabled robust quantitative analysis and visualization of HER2 motion in protrusion and non-protrusion subcellular regions of breast cancer cells. We report that signaling of protrusion growth is a local event that is transduced by unimpeded, rapid HER2 spatiotemporal dynamics in filpodia. These results indicate the value of emerging efforts focused on better controlling cancer signaling pathways via filopodia and other protrusion-targeted therapeutics.

## Results

### Segregated, Distant Morphology of Breast Cancer Filopodia and Lamellipodia, in Vitro and in Vivo

Filpodial and lammelipodial protrusions are present at the invasive front of cancer cells *in vitro*, and in *in vivo* tumor spheroid models, but their morphology - particularly in *in vivo* tumor tissue is less studied [10, 35], We used light and electron (SEM/FIB-SEM) microscopy to examine protrusion organization in *in vitro* cell models and asked if this organization was similar in *in vivo* breast tumor tissue.

We first examined the morphology of SKBR3 cells, a HER2-amplified epithelial breast cancer model used for investigating HER2 and cancer protrusion biology (e.g [18, 19]). SKBR3 protrusions are composed of finger-like filopodia that extend from sheet-like lamellipodia and lie in protrusion-rich regions at the polar ends of the cell body (**Figure 1A)**; they contact the substrate surface and participate in cellular motility (**Figure 1B, Movie 1**). A striking feature of filopodia is their thin diameter, relative to their remote distance from the cell body (**Figure 1B, Movie 2).** Filopdial widths measure 80-130 nm in diameter but their distance from the cell body is 2-5µm (from the lamellipodial base to filopodial tip) (**Figure 1C**). This was also evident in HER2-amplified 21MT-1 epithelial and low HER2-expressing MCF7 luminal breast cancer cell models (**SFigure S1**). Protrusions also appeared as morphologically distinct organelles, almost separated from the cell body; in instances we observed filopodial-lammelipoidial protrusions connected just peripherally to the cell body through a few membrane extensions (**Figure 1B, last frame)**.

**Figure 1.**
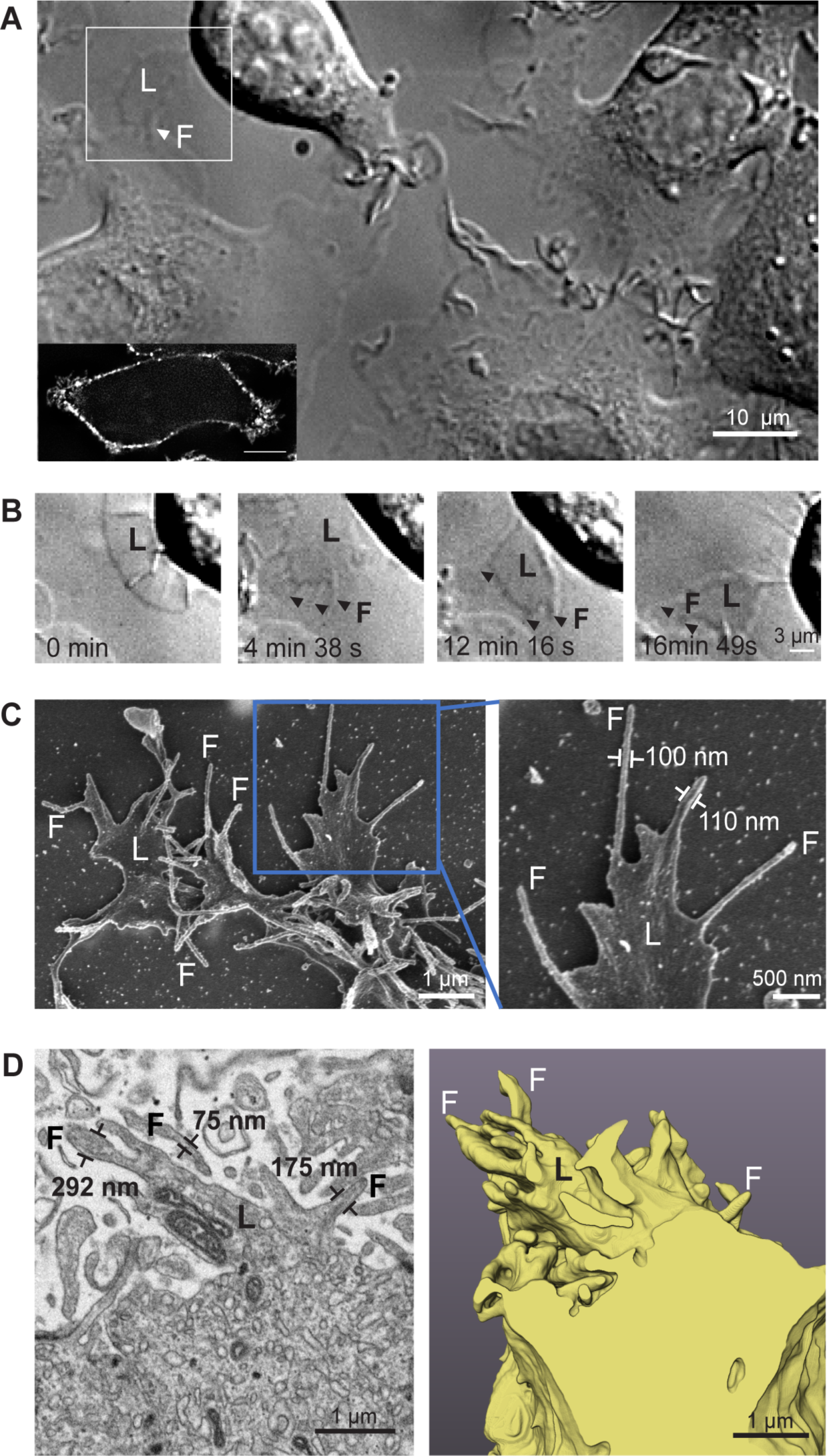
Segregrated morphology of breast cancer cell protrusions, *in vitro* and *in vivo*. **(A)** Movie frame shows a motile, SKBR3 breast cancer cell, in serum media, *in vitro*, extending protrusions (see **Movie 1**). Example lamellipodium (L), filopodium (F). Inset: wheat germ agglutinin-Alexa488-membrane labels shows dense protrusion and non-protrusion regions of the SKBR3 cell body; scale bar 2um. **(B)** Frame stills (*white box*, Figure 1A) shows morphologically segregated nature of lamellipodia and filopodia. A lamellipodium (L, 0 sec) extends numerous filopodia (F, 4 min 38 sec) which is peripherally situated and, at times, almost separated from the cell body (F, 16 min 38 sec); (see **Movie 2**). **(C)** SEM shows slender filopodia (F) measuring ∼ 100 nm wide, ∼1um long that extend from lamellipodia (L) which measure additional distances of several microns further from the cell body; SKBR3 cells *in vitro*. Additional human breast cancer cell lines shown in **Supplement Figure S1. (D)** Focused-ion beam-EM shows similar segregated, far-reaching filopodial and lamellipodial morphological organization in human *in vivo* HER2-positive breast tumor. *Left:* cell from a biopsy of a metastatic bone lesion imaged by volume EM (single 5 nm slice). *Right:* segmented, 3D, rendered volume of the lamellipodium (L) and filopodia (F). See **Movie 3** for volumetric stack overlay and **Supplement Figure S2** for wide-field view of stained resin section of the tumor biopsy.

To determine if these morphological features were also present in intact tumor tissue, we used 5nm resolution FIB-SEM to image biopsies taken from a HER2+ metastatic breast tumor. FIB-SEM images revealed nanoscale protrusions at the polarized apical surfaces of cells that extended into the extracellular tissue matrix (**Figure 1D**, *left*). A complete 3D reconstruction of a protrusion showed the *in vivo* architecture of a filopodia extending from the body of a lamellipodium (**Figure 1D**, *right*; **Movie 3**). The lamellipodium was widest, 1.3µm, at the base where it joined the cell and tapered in width to 690nm where the filopodia began their extension. Filopodia contained some ribosomes but had very few large organelles compared to lamellipodia which contained mitochondria, vesicles, and ribosomes; no junctional complexes were observed at these contact sites. Together, the lamellipodium and filopodia extended into an intercellular space and had sites of membrane to membrane contact with filopodia from neighboring breast cancer cells. The width of each filopodium varied along its length, with the average diameter of filopodium extending from the lamellipodium measuring 140 nm. The length of the complete structure, from where the lamellipodium joined the cell to the tip of the longest filopodia, was 3µm. These data show the remote, segregated nature of nanoscale breast cancer filopodia that is characteristic of both *in vitro* and *in vivo* cells and tissue.

### HER2-PI3K Signaling is Protrusion-Compartmentalized

To identify the spatial locales of HER2 signaling, we measured the local concentration of HER2 and pAKT complexes in protrusions and the cell body of SKBR3 breast cancer cells. HER2 and its PI3K pathway surrogate, pAKT, are associated with breast cancer cell protrusions [18, 19, 36]; however, these levels of HER2 and HER2-evoked PI3K signaling have not been established quantitiatvely and in relation to the rest of the cell.

STORM imaging of HER2-stimulated, herceptin-Alexa647-labelled SKBR3 cells revealed discrete HER2 complexes present in both finger-like filopodia and fan-shaped lamellipodia; HER2 complexes were not detectible in the rest of the cell (**Figure 2A**). Neuregulin 1 (NRG1)/ heregulin is a growth factor that induces the formation of HER2-HER3 heterodimers and evokes PI3K pathway-mediated formation of protrusions [37, 38]. Fluorescent imaging of NRG1-stimulated, anti-HER2 affibody-quantum dot (QD)-labelled probes showed discrete protrusion-located HER2 complexes in SKBR3 cells. The HER2 affibody-QD probes were custom designed and generated by monovalently linking a small recombinant protein consisting of a HER2 affibody (58 amino acid peptide) Z_HER2:342_ fused to a fluorogen activating protein (FAP_dL5**_) to a QD (**Figure 2B, Methods, Figure S4**).

**Figure 2.**
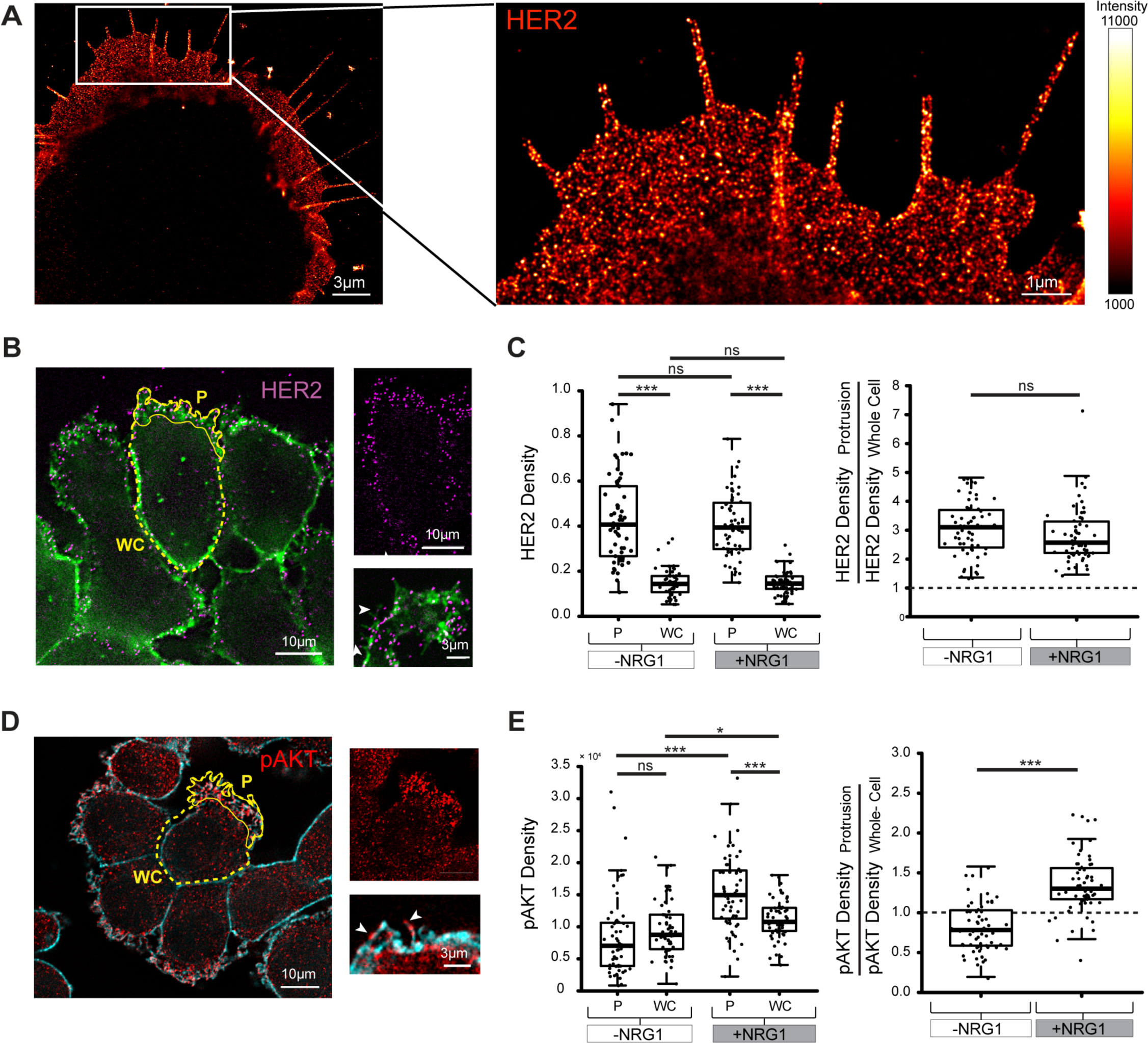
Protrusion-Contained HER2-PI3K Signaling. **(A)** HER2 localization in filopodia and lamellipodia shown by STORM imaging. Herceptin-Alexa647, SKBR3 cells in serum. **(B)** Localization of HER2 in filopodia and lamellipodia shown by discrete HER2 affibody-quantum dot labelling (*magenta*); quantitated in C). +NRG1, SKBR3 cells. Annotated protrusion regions (*P, solid yellow line*) distinguished from rest of whole cell in same z-plane (*WC, dashed yellow line*) by wheat germ agluttinin-labelled plasma membrane (green). Right hand side: *(top)* HER2 channel alone showing discrete HER2-QD complexes; *(bottom)* zoomed image showing HER2 on filopodia (*white arrows*) and lamellipodia. **(C)** Plots showing higher HER2 density at protrusion vs. whole cell regions;HER2 density remains at same levels in protrusions before and following NRG1 activation. **(D)** pAKT immunolabeling shows higher density of pAKT label at protrusions, quantitiated in E). Right hand panels: *(top)* pAKT channel alone and *(bottom)* zoomed detail of pAKT complexes on filopodia (*white arrows*) and lamellipodia. **(E)** Plots showing higher pAKT density at protrusion vs. whole cell regions; pAKT levels significantly increase following NRG1+ activation and are predominantly localized at protrusions. In **(C)** and **(E)**, HER2 density computed as number of HER-QD complexes and pAKT density computed as total pAKT-Alexa488 fluroescence, per unit area of P and WC regions at z-plane of protrusions (*left*) and as a ratio of HER2 density in protrusion:whole cell regions (*right*). +NRG1 is 15 mins stimulation in serum-free media, -NRG1 is serum-free. *** p<0.001, * p<0.05, ns= non-significant, Mann-Whitney rank sum test. Each dot is an individual SKBR3 cell collected from three different experiments, n=118-119 cells. See **Supplementary Information 9**, statistics.

To quantitate the protrusion-localized nature of HER2 as well as pAKT, we computed the spatial density of HER2 and pAKT at protrusions at the plane of the glass, where protrusions are found. Spatial densities of HER2 in protrusion regions and for the rest of the cell were obtained by counting the number of HER2-labelled QD complexes and dividing this count by the area of the entire cellular region at the same z-plane as protrusions (where protrusions make contact with the substrate surface). Cellular regions were annotated using fluorescently-labeled WGA, a lectin that binds to sialic acid and *N*-acetylglucosaminyl residues on the plasma membrane (**Figure 2B**: *P, EC*). The spatial density of HER2-QDs in protrusions is 2-3 times higher, on average, compared to the total cell area (**Figure 2C**, *left plot*). Moreover, HER2 density in protrusions remains steady and is indistinguishable prior to and following NRG1 stimuluation. Additional experiments showed HER2-QDs were not found in the volume of the cytoplasm of the cell body even at 60 minutes following NRG1 stimulation; (**SFigure S4**). Thus, the level of HER2 is highly enriched in protrusions and remains steady in protrusions following HER2 activation (**Figure 2C**, *plots*). pAKT labelling under the same physiological conditions revealed discrete pAKT-Alexa647 complexes that were visible along filopodia, in lamellipodia and cell soma (**Figure 2D).** The spatial density of pAKT-Alexa647 fluorescence showed that baseline levels of pAKT at protrusions were indistinguishable from levels computed for total cellular area, at the same z-plane (**Figure 2E**, *left*). However, following NRG1-evoked HER2 signaling pAKT activation was significantly enriched at protrusions compared to the entire cell area (**Figure 2E**, *right*), indicating that downstream PI3K-signaling that is evoked by HER2 remains spatially localized within protrusions. These quantitative data show that activation and downstream propagation of HER2 signaling is a protrusion-localized event that is separate from the main cell body.

### HER2-Evoked Protrusion Growth Output is Proximal to HER2-Activation Sites

To identify the spatial extent of HER2 signaling outputs, we examined the location of new protrusion formation in proximity to sites of HER2 stimulation. Global, bath-applied NRG1 stimulation produces cell-surface ruffling and the formation of protrusions at the substrate surface in SKBR3 cells (**Figure 3A; Figure S4)**. We used a glass micropipette filled with NRG1 to evoke subcellular localized HER2 signaling (**Movie 4)** and found that SKBR3 cells responded rapidly, producing new protrusions within 3 minutes after stimulation, at sites proximal to the location of NRG1 delivery (**Figure 3B**, *first row* and **Movie 5**). Localized protrusion extension could be evoked in adjacent cells and was absent in unstimulated portions of the cell (**Figure 3B**, *second row*; **Movie 6**). Furthermore, control experiments showed using a micropipette filled with phosphate buffered saline did not evoke protrusion formation (**Figure 3B**, *third row*). siRNA silencing of HER2 expression confirmed that the NRG1-evoked signaling was mediated by HER2 receptor inputs: 1) decreased HER2 expression was measured by Western blotting and molecular counting of discrete HER2 complexes and 2) reduced HER2 expression decreased the fraction of cells exhibiting filopodia (**Figure 3C**). Thus, HER2-evoked downstream signaling, from initiation of HER2 activation to evoked new protrusion growth is rapidly responsive, protrusion-localized process.

**Figure 3.**
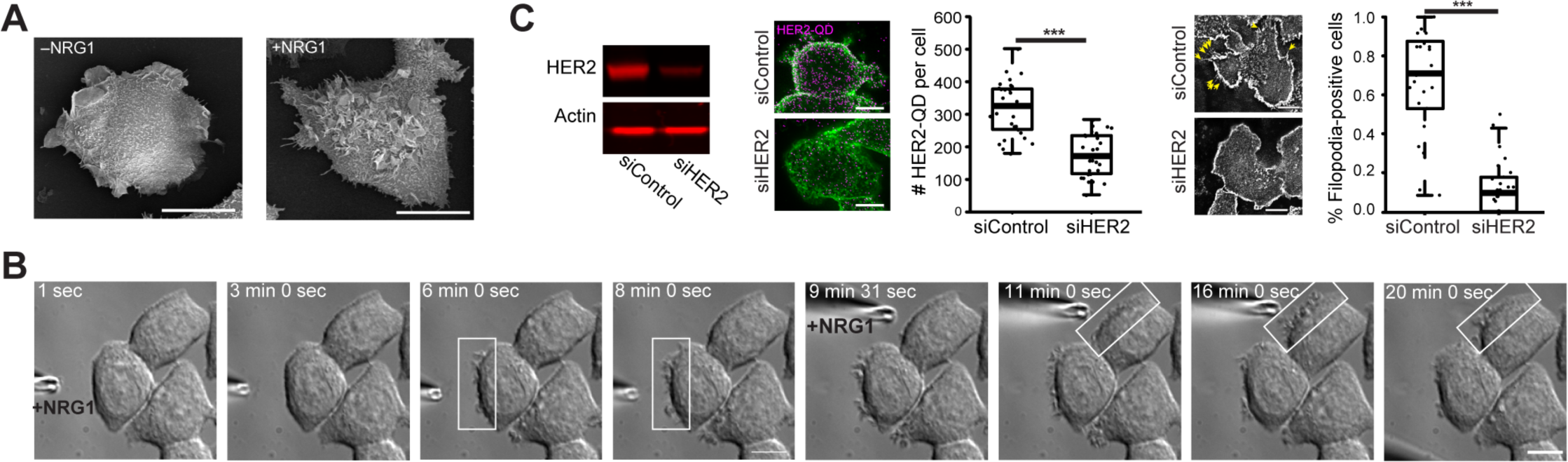
New Protrusions Growth Evoked Near the Site of HER2-PI3K Pathway Stimulation. **(A)** NRG1-stimulated (15 minutes) SKBR3 cells exhibit membrane ruffling and increased surface-attached lamellipodia and filopodia compared to control cells, shown by SEM. **(B)** Subcellular-localized delivery of NRG1 via micropipette to two different SKBR3 cells shows rapid new protrusion growth within < 3 mins (frames from **Movie 5** & **6)**. Control micropipette containing PBS does not evoke protrusion formation (**Supplement Figure S5**). **(C)** HER2 siRNA reduces HER2 expression and filopodial growth compared to control scrambled siRNA-treated cells, shown by qualitative Western blot; immunolabelling of HER2 with HER2affibod-QDs with quantiation of discrete HER2-QD counts per slice per cell; and quantification of filopodia-positive cells. Cells stimulated with 15 mins of NRG1 in both HER2 siRNA and control siRNA conditions. % filopodia-positive cells quantified from the WGA-membrane labeled fluorescence channel of same experiments as immunofluorescence experiments. Example monochrome images of WGA-labelled cells show filopodia (*yellow arrows*). For plots, *** p<0.001, Mann-Whitney rank sum test conducted with data from three different experiments, 30 ROIs, n=721 cells. See **Supplementary Information 9**, statistics. All scale bars=10um.

### HER2 Spatiotemporal Dynamics Differ in Filopodia vs. Lamellipodia and Non-protrusion Regions

To understand the HER2 spatiotemporal dynamics underlying protrusion signaling, we performed single particle tracking of HER2 complexes in live SKBR3 cells. Using compact, monovalent HER2 affibody-QD probes, we observed individual HER2-QD complexes for extended time periods (2-4 minutes) (**Methods, SFigure S4**).

NRG1 stimulation increased HER2 mobility and speed in cells (**SFigure S6**) and HER2-QDs were found in dynamic motion in filopodia, lamellipodia, and plasma membrane regions absent of obvious protrusions (F, L, NP, **Figure 4A)**. HER2-QDs also were found moving on protrusions on the under-side of the cell (‘S’, **Figure 4A; Methods)**; these dynamics were not studied because of potential interactions with the slide surface.

**Figure 4.**
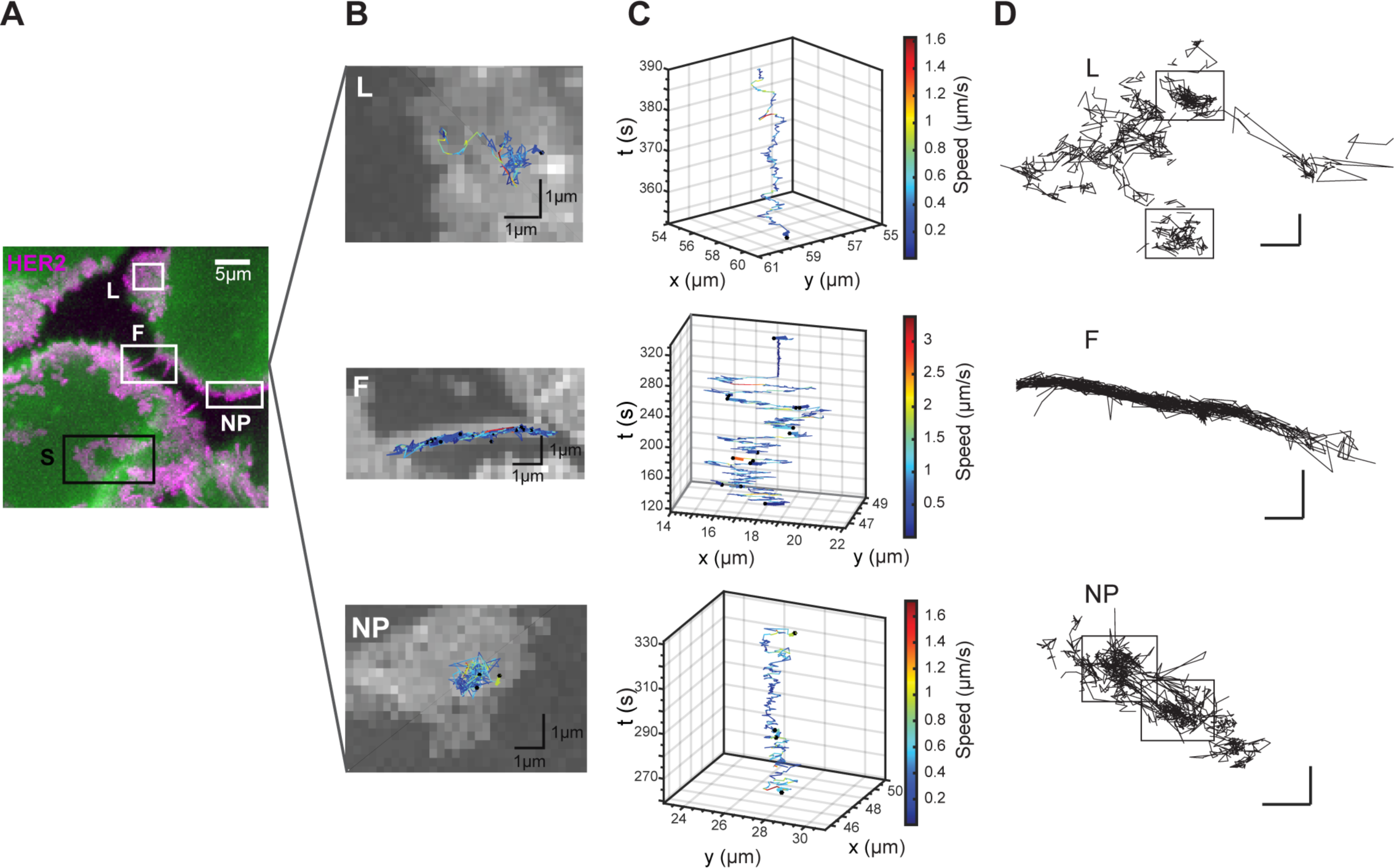
Features of Filopodial HER2 Spatiotemporal Dynamics that are Distinct from Lamellipodia and Non-protrusion Subcellular Regions. **(A)** HER2 trajectories in subcellular regions of filopodia (F), lamellipodia (L), no protrusion subcellular (NP), and under-side surface of cell in contact with the glass substrate (S), shown by maximum intensity projection of HER2-QD fluoresecence over time in SKBR3 cells cultured in serum-media. (magenta, HER2 affibody-QD probe, green, WGA-labelled plasma membrane). **(B**, **C, D)** Example individual HER2-QD trajectories showing in filopodia, the rapid speed and back-and-forth linear motion compared to slower, non-linear ‘picket-fence’ motion lamellipodia and cell body. Overlay of trajectories color-coded to denote instantaneous velocity on max intensity projection of HER2-QD over time (B) and by *x-y-t* plots (C). **(D)** additional HER2-QD trajectories showing zoomed detail of compartmented ‘picket-fence’ motion in lamellipodia and non-protrusion regions. See **Movie 8-13** for corresponding videos. All scale bars= 500 nm.

HER2 dynamics differed depending on their subcellular locale. In filopodia, HER2-QDs moved in a linear fashion along the filopodia longitudinal axis; this motion was notably rapid. Instantaneous speeds of (>1 µm/s) were found along filopodia and reached even up to (2-3 µm/s) (**Figure 4B**), values which lie in the upper range of speeds reported for active transport of receptors (1-2 um/s) **[23, 39-41]**. At the filopodia tip and base, HER2-QDs moved in a back-and-forth fashion or stayed static before continuing motion along the length of filpodia (**Figure 4B, Movie 9-10**). Her2 motion in lamellipodia and non-protrusion regions appeared markedly different from filopodial motion. HER2 motion in both lamellipodial and non-protrusion regions appeared randomly-oriented and HER2-QDs moved at slower instaneous speeds (0.2 −0.6 um/s) with maximum speeds < 1.6 um/s (**Figure 4B**; **Movie 11-12** (L), **13-14** (NP)). Moreover, HER2 motion in lamellipodia and non-protrusion regions appeared temporarily hindered at times by barriers which corralled HER2 motion into regions spanning a few hundred nanometers (**Figure 4C)**. These data reveal the rapid, linear motion of HER2 in filopodia that contrasts slower, corralled motion in lamellipodia and non-protrusion regions.

### Unimpeded Brownian Mechanics Underlies Filopodial HER2 Rapid Dynamics

To better understand the molecular basis of the HER2 motions we observed, we analyzed HER2-QD motion in each of the subcellular regions. Active, cytoskeletal motors are one form of direct, linear transport that has features of the motion we see in filopodia, and free diffusion that is hindered by physical barriers such as cytoskeletal corrals, qualitatively resembles the motion we see in lamellipodia and non-portrusion membrane regions [42]. Both these forms of motion are proposed to underlie membrane protein trafficking and signaling in cilia and other types of protrusions [42-44]. We tested the hypothesis that active transport is a main mechanism of HER2 motion in filopodia, whereas hindered free diffusion underlies HER2 motion in lamllipodia and non-protrusion portions of the membrane in breast cancer cells.

We used standard diffusion analysis to quantify the diffusion-like mobility of HER2. The MSD data show that HER2 mobility, quantified by its effective diffusion constant, is ∼3-4 times higher in filopodia than in lamellipodia and non-protrusion plasma membrane regions (**Figure 5A, Figure S7A)**. Despite clear differences in the magnitudes of the MSDs for the different cellular regions, none of the regions exhibited simple linear behavior (**Figures 5A**,**B; SFigure 7**) motivating additional analysis. We probed whether HER2-QD motion in filopodia resulted from passive or active motion, based on two analyses. To test the hypothesis that HER2 filopodial motion resulted from passive simple diffusion confined by insurmountable barriers, we simulated confined simple Brownian diffusion (**Methods**) and found the MSD behavior to be indistinguishable from that observed for HER2-QD in filopodia and that the slope of the experimentally observed MSDs never exceed a value of one on a log-log plot (**Figure 5B**, **Figure S7B)**. Secondly, to assay the presence of driven motion more directly, we performed an autocorrelation analysis (**Methods**). The results show that at no point along the length of filopodia was there positive correlation - a tendency to continue stepping in the same direction as the prior step (**Figure 5C**); positive autocorrelation (see Figure S7C) would be expected for directed, motor-driven motion. In addition, experimental filopodia data again are indistinguishable from data obtained from passive, (unconfined and confined) Brownian simulation. The simulation comparison and autocorrelation analysis confirm the HER2-QD motion in filopodia is passive in nature.

**Figure 5.**
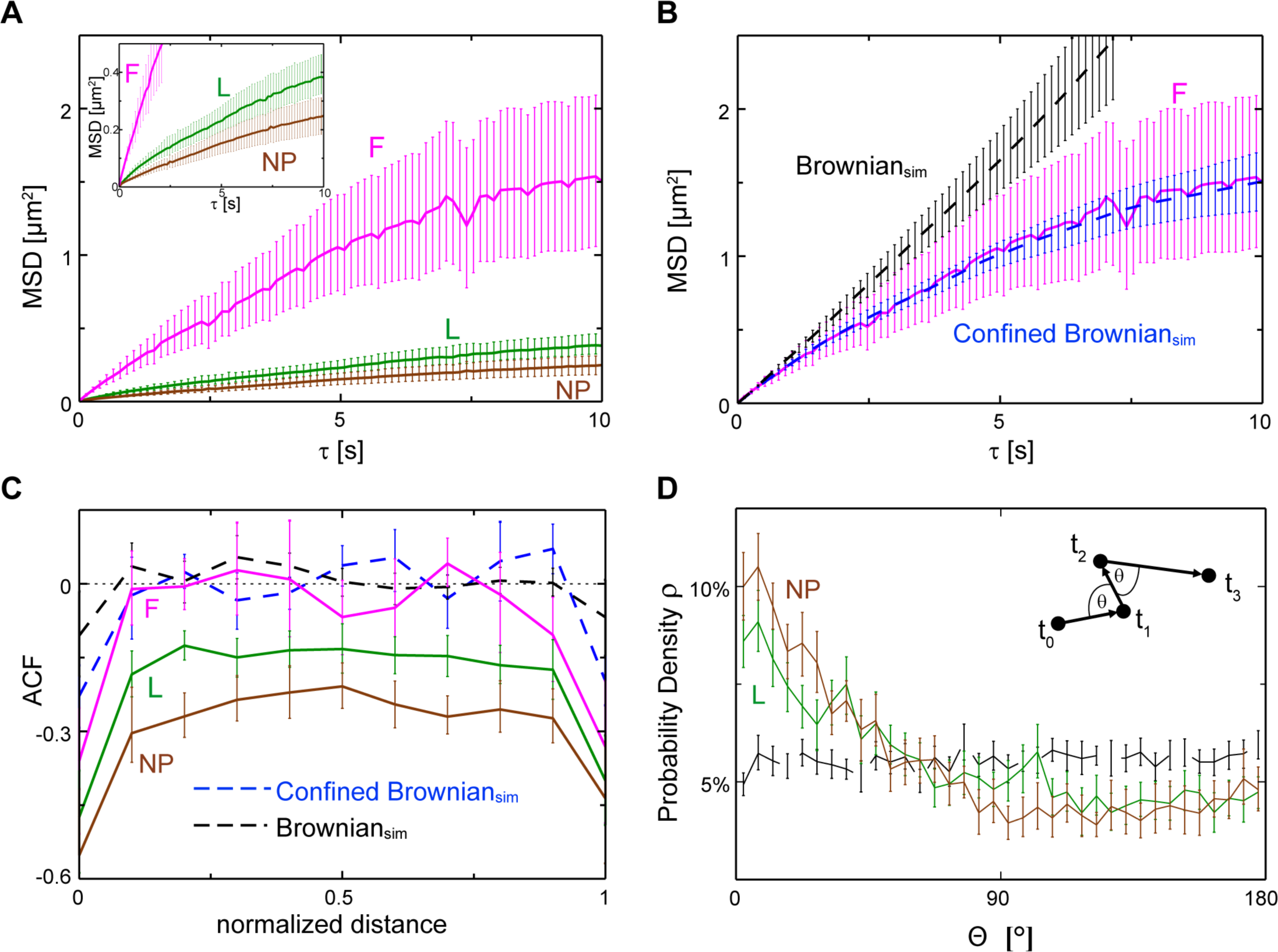
Diffusion and correlation analysis of locus-specific HER2 motion. **(A)** MSD is plotted as a function of the time interval τ for HER2-QD mobility in the filopodia (magenta), lamellipodia (green), and the non-protrusion (brown) regions of the cell. The inset in the upper left corner is a zoomed-in version with MSD < 0.5 µm^2^. **(B)** In a comparison of simulation with experiment, MSD of the 1D confined (blue dashed line) and unconfined 2D simulations (black dashed line) are shown along with the filopodial MSD (F, magenta) which precisely matches confined simulation. **(C)** Two-step auto-correlations (ACFs) are plotted as a function of the normalized distance traversed in the different regions and simulations (same colors and line styles as in (A) and (B)). An ACF of zero indicates directionally uncorrelated motion while negative values indicate the presence of direction reversals. Note that the ACF is negative throughout the lamellipodia and the non-protrusion regions. **(D)** The distribution of the angle, Θ, between two subsequent steps for the lamellipodia (green) and the non-protrusion (brown) regions as well as for the unconfined 2D simulation (black dashed line). The higher probability of small angles in these experiments indicates the frequent occurrence of reflections (see inset). In (A) and (B), the mean squared displacement was computed for the 1D motion of HER2-QDs along the length of filopodia and the 2D motion of HER2-QDs in lamellipodia and protrusion-free cell regions. All MSD data were shown are projections to a single dimension to enable direct comparison (**Methods**).

The behavior in the non-filopodia regions, lamellipodia (L) and non-protrusion (NP) regions, were subjected to an analogous characterization. In the distribution of step angles (**Figure 5D**), the over-representation of small-angle displacements compared to truly random simple diffusion directly shows that HER2-QD trajectories in lamellopodia and the non-protrusion regions experience frequent reflection events. This confirms the tendency for step reversals revealed by the negative autocorrelation (**Figure 5C**). The always-negative autocorrelation profiles of L and NP regions (**Figure 5C**) suggest that reflections occur (i) at the interior of regions explored by individual trajectories and therefore (ii) at *surmountable* barriers, consistent with direct inspection of 2D trajectories in lamellipodia and cell body which show that HER2-QD mobility is occasionally impeded, but not confined, by some type of sub-micron structure (**Figure 4C**). Hence, in contrast to the filopodial behavior, HER2-QD motion in lamellipodia and non-protrusion regions exhibited distinct characteristics consistent with a ‘picket-fence’ model of surmountable membrane compartments, which is further highlighted by the non-Gaussian distribution of HER2 step sizes in these regions (**Figure 7D**,**E**).

To relate the HER2-QD temporal measurements to subcellular regions, we mapped the positional density and speed of individual HER2-QD trajectories. In filopodia, the distribution of HER2-QD positions are uniform (blue squares, **Figure 6A**) whereas lamellipodia and non-protrusion regions contained regions over which the HER2-QD traversed repeately (yellow-red squares, **Figure 6B, 6C**). Velocity vectors show that nanoscale motion of HER2-QD along the length of filopodia (long vector lengths) is fast with reversals at the tip and base of filopodia (**Figure 6B**, filopodial tip; **SFigure 6A**, filopodial base). In contrast, L and NP regions show reflected and slowed motion (short vector lengths) of HER2-QDs which are temporarily corralled by a sub-micron compartment (**Figure 6B, 6C, SFigure 6B**) and is consistent with the temporal analyses (**Figure 5A, 5C, 5D**).

**Figure 6.**
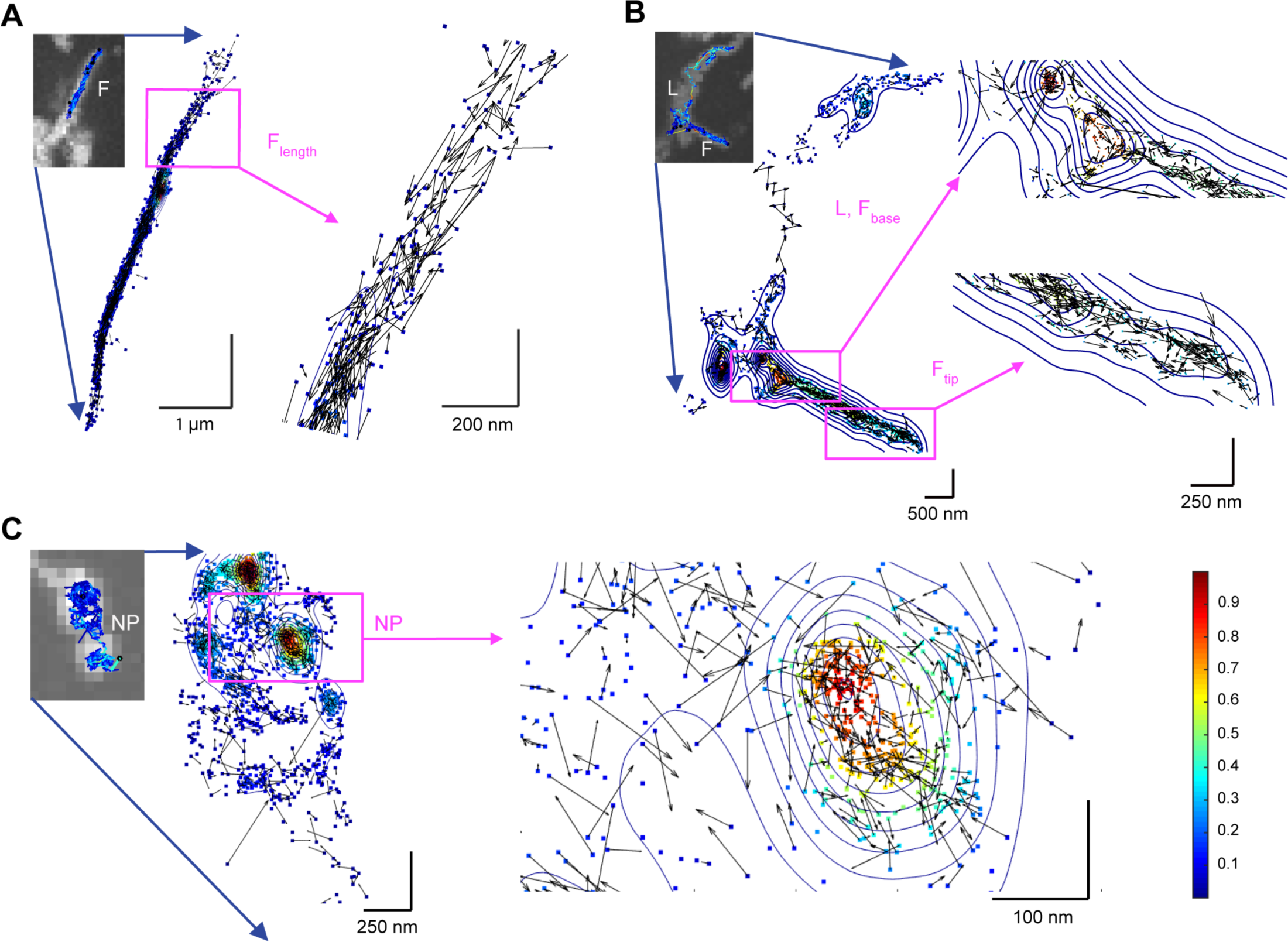
Mapping of HER2 Positional Density and Velocity by Subcellular Region. Positional densities are shown as color-coded squares and contour lines and normalized for each trajectory (see colored scale bar). Velocity vectors are denoted by arrows with component (v_x_, v_y_) at the points (x, y). **(A)** Representative individual HER2-QD trajectory in filopodia showing uniform distribution of HER2 motion with direction along the filopodial longitudinal axis. **(B)** Representative individual HER2-QD trajectory showing regions over which the HER2-QDs repeatedly traverses the lamellipodia (L) and the base of filopodia (F_base_). Contour lines show the circular shape of these surmountable, sub-micron compartments in which velocity vectors show reflected motion toward the center. At the filopodial tip (F_tip_), HER2-QDs encounter reversal in direction. **(C)** Representative individual HER2-QD trajectory in non-protrusion region showing circular, surmountable sub-micron compartments at which HER2-QDs show reflected motion toward the center. See **Figure S8** for additional examples.

Together, these data reveal a rapid mobility of HER2 in filopodia that does not arise from active, directional motors but instead, closely approximates passive Brownian motion along the length of filpodia, confined by the base and tips of filopodia. This filopodial motion differs from compartmented “picket fence” motion we observed for HER2 in lamellipodia and non-protrusion regions of the same cells.

### Modulation of Linear Actin Reduces HER2 Mobility and Downstream PI3K-pAKT Signaling

Early, growing studies using blockers of actin and other cytoskeletal filaments, report evidence pointing to the role of actin and other cytoskeletal nanostructures in modulating the receptor speed and its consequent signaling in immune and neuronal cells [45, 46]. Given the rapid speed of HER2 in filopodia and that filamentous actin is a major component of filopodia, we sought to understand impact of HER2 mobility on HER2-PI3K signaling in by modulating actin structure in filopodia using latrunculin B (LatB) block of actin polymerization [47, 48] and fascin inhibitors, which specifically target filopodial actin assembly in actin-rich cancer cell filopdia and reduce cell invasion [49].

2D structured illumination microscopy shows linear actin closely-apposed to the plasma membrane of protrusive triangular-shaped lamellipodia that converges into filopodia (**Figure 7A**, *left top*). LatB-treatment led to the reversible retraction of filopodia with reduced levels of actin (**Figure 7A**, left bottom). Upon LatB removal, cells re-extended protrusions, although not to the same length (Movie 15). As expected, Lat B disrupted the linear geometry of filamentous actin fibers; 3D STORM imaging shows LatB+ retracted protrusions appearing curved and flattened (**Figure 7A**, *right, top and bottom;* **Movie 16-17**). HER2-QD particle tracking showed that LatB+ block of actin polymerization did not stop HER2 motion but reduced its speed and mobility (top plots, **Figure 7B**, top; **Movie 18-21**). This slowed HER2-QD mobility resembled NP regions of cells in LatB-cells (red and brown curves, **Figure 7B**, *bottom*); the mean squared displacement (MSD) of HER2-QDs under LatB treatment was not distinguishable from HER2-QD motion measured in NP regions (**Figure 7B**, *bottom*). Furthermore, LatB+ cells showed decreased pAKT signaling as confirmed by quantitative measurement; NRG1 stimulation produced a significant increase in pAKT at the protrusion but was indistinguishable from the rest of the cell with LatB+ NRG1 stimulation (left, **Figure 7C**)). The decrease of pAKT signaling in protrusions was also evident with cells treated with the inhibitor of filmentous, linear actin found in filopodia, G-Fascin (*right*, **Figure 7C**).

**Figure 7.**
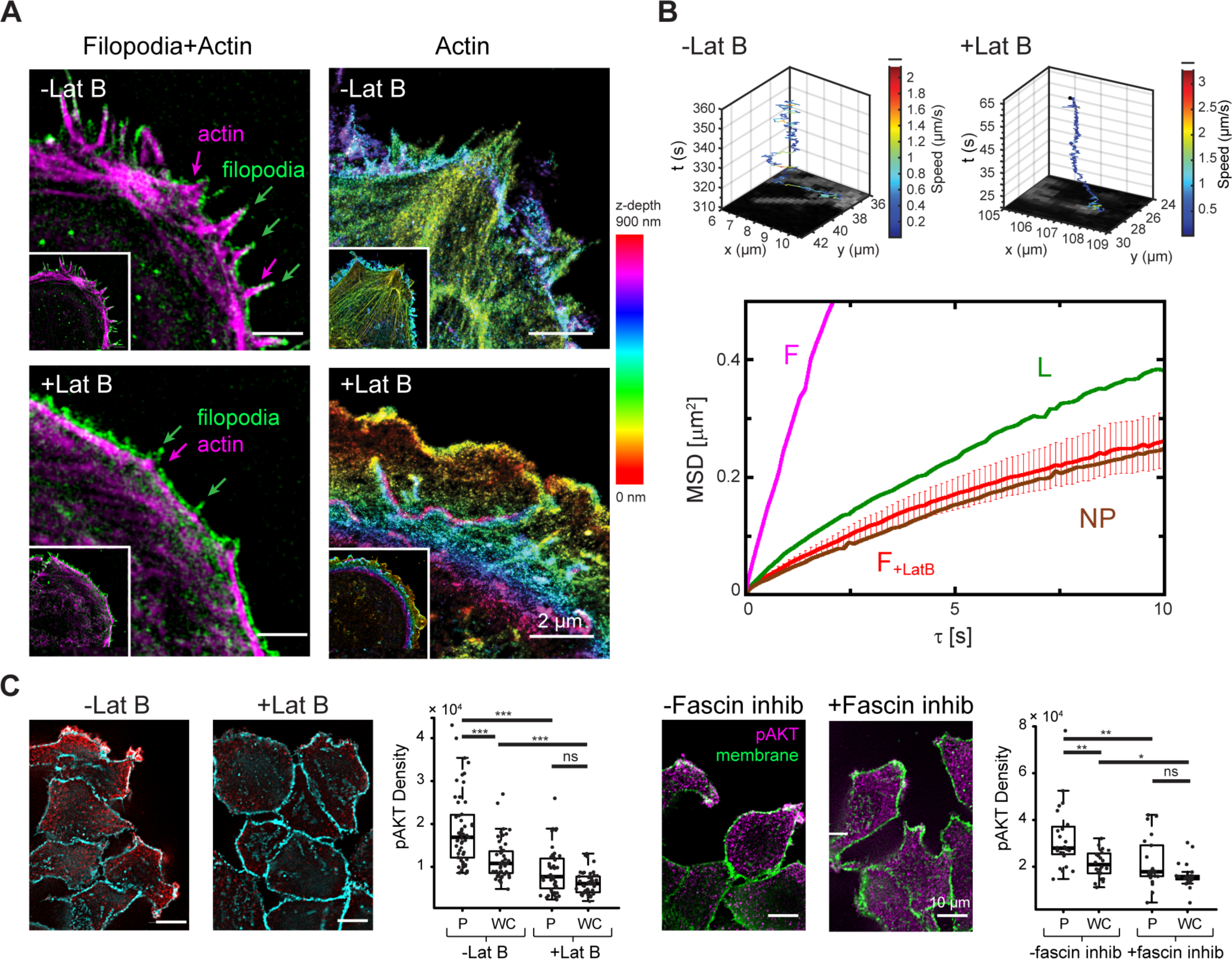
Reduced HER2 mobility Reduces pAKT downstream protrusion signaling. **(A)** Trajectories showing HER2-QD mobility latrunculin-treated (LatB+) SKBR3 cells HER2-QD mobility measured at the peripheral plasma membrane in latrunculin-treated (LatB+) SKBR3 cells shows slowed mobility compared to (LatB-) **(B)** HER2-QD mobility in filopodia (left plot). HER2-QD mobility under LatB+ is similar to HER2-QD motion in LatB= lammelipodial and in protrusion-free portions of the plasma membrane (L, NP). Filopodial protrusions contain actin with linear structure that is perturbed following LatB treatment (+LatB). 3D super-resolution of actin in SKBR3 cells cultured in serum-media. Colored bar, axial depth. See 3D rendering in **Movie 17, 18**. HER2-QD trajectory on filopodium of -LatB and on altered protrusion stubs of +LatB of SKBR3. Colored bar, HER2 speed. See **Movie 19-22. (D)** HER2-QDs on +LatB altered protrusions show slower mobility that resembles HER2-QD motion on cell body. Cumulative probability distribution of HER2 squared displacement. *** p<0.001, KS test. **(E) (G)** HER2-QD (magenta) is localized on actin-rich filopodial protrusions (*arrowhead*) in SKBR3 cells in serum-media. **(H)** pAKT-Alexa488 (magenta) is also localized on actin-rich filopodial protrusions (*arrowhead*) in SKBR3 cells stimulated with NRG1. **(I)** pAKT, concentrated in protrusion-rich regions (*arrowhead*) after NRG1 stimulation (-LatB), is reduced after latrunculin B (+LatB). pAKT-Alexa488 (red) and WGA-Alexa647 membrane (cyan). +LatB: 15 mins NRG1 and LatB in serum-free media; -LatB: 15 mins NRG1 alone in serum-free media. Values in **(D), (E)** from cellular regions: CB, L, and F are replotted from **Figure 4C, 4D** for comparison to +LatB data. n=276 receptors, 95 cells. See **Supplementary File 1**, statistics.

Together, these data show that disruption of linear, filamentous actin geometry in filopodia does not stop HER2 dynamics but reduces its mobility to a degree that resembles that of non-protrusive regions of the cell and diminishes HER2-PI3K downstream signaling activity in protrusions.

## Discussion

Filopodial and lammelipodial protrusions are thin, fine celluluar structures that are increasingly recognized as key players in the detrimental processes of cancer cell invasion and tumor spread, the primary cause of cancer deaths. Here, our studies show that HER2 signaling of breast cancer cell protrusion formation is conducted in filopodial and lammelipodial nanoscale organelles in a site-localized manner, remote from the cell body: 1) HER2 molecules are present in protrusion structures and absent from the rest of the cell, 2) HER2 remains sequestered at protrusions regardless of HER2 activation state, 3) HER2-evoked pAKT-PI3K activation is protrusion-localized, and 4) formation of new protrusions is rapid and proximal to the site of HER2 stimulation. This signaling is conducted in part by HER2 mobility in filopodia that is particularly rapid and unexpectedly does not follow commonly expected ‘picket fence’ barrier model of surmountable compartments in the plasma membrane, nor is it directed, active cytoskeletal transport. Instead, filopodial HER2 dynamics resemble unimpeded Brownian motion that enable HER2 to move with 2-3 times higher mobility than in lamellipodia/non-protrusion regions of cells, and at speeds (2-3 um/s) that are equivalent to active motor transport requiring active energy sources.

While HER2 and pAKT have been observed at breast cancer cell protrusions and mechanisms of HER2 regulation have been proposed [19, 20, 22], we provide first-time evidence here, through quantitative measurement, that HER2 activation and its evoked pAKT-PI3K outputs when compared to the rest of the cell, is protrusion-centric **(Figure 2)** that importantly results in functional formation of new protrusion growth near the vicinity of HER2 activation **(Figure 3)**. Furthermore, this protrusion-centric nature of functional HER2-PI3K protrusion formation is aligned with the morphologically-segregated and remote nature of filopodia and lamellipodia that we observe, *in vitro* and *in vivo* (**Figure 1**). Given the importance of protrusion abundance in tumor invasion and progression, these combined data, provide strong indication for emerging efforts in developing therapies that target the protrusion as means to inhibit protrusion growth. Indeed, mutations in proteins specific to filopodia have been recently identified which could serve as potential effective targets; our results showing block of actin in protrusions and specifically in filopodial decreased HER2 mobility and pAKT signaling (**Figure 7**) suggesting that compounds that modulate the integrity of the filopodial environment components such as cytoskeletal actin could as well produce means of improved, potent control [7, 9, 10, 50]. Such protrusion-directed therapies could be used alone or in combination with small molecule inhibitors that directly target signaling pathway molecules to control tumor growth but which unfortunately often lack enduring control of tumor growth.

Our results showing that filopodial HER2 motion is fast, lacks membrane ‘picket fence’ barriers, and that this fast transport is conducted not by active motor transport but passive Brownian mechanics are unexpected. Filopodial HER2 filopodial speeds are equal to active motor-driven transport of receptors measured in live cells (1-2 um/s) and yet are not driven by active transport but passive Brownian motion (**Figure 6**). Moreover, filopodial HER2 mobilities are not only x2-3 faster than HER2 in lamellipodial and non-protrusion cell regions but also for general classes of membrane receptors in other cell systems, measured by single particle tracking (**Supplementary Table 1**). Lammelipodial HER2 and non-protrusion HER2 mobilities show temporary membrane compartmented barriers that are expected for protein motion in the membrane; the unusual finding that filopodial HER2 dyanamics are unimpeded – except at the base and tip of filopodial-may explain in part how these rapid dynamics are possible. Recent studies have reported observations of actin in modulating receptor mobility, and have proposed models in which ‘linear channels’ and ‘microdomains’ defined by actin regulate these aspects in immune cells [45, 46, 51]; it is very possible that we observe a similar phenomenon in which rapid dynamics are made possible through linear conduits of actin in filopodia-while toxin block of actin polymerization has experimental limits – it is notable that LatB and fascin inhibition of filopoidial actin produces a mobility of HER2 that is not blocked but resembles HER2 motion in non-protrusion regions of the same cells. While there could be a high-cost for the simple design of the filopodia, it appears that filopodia may be designed to conduct rapid HER2 signaling in an energy-efficient manner.

What is the function of segregating signaling molecules to the protrusion organelle? While there are many examples of locally sequestered molecules, presumably to strengthen signaling (e.g. membrane rafts, cytoskeletal corrals, signaling endosomes, synaptic densities), it is notable that such localized cell mechanisms eventually propagate these signals to other cellular sites (e.g. from plasma membrane to cytosol, from periphery of cell toward the nucleus). Interestingly, while this contained signaling segregation at breast cancer cell protrusions is unusual, it is not singular-in the nervous system, where fine dendrites and axons of neurons can span far distances of millimeters from the main cell body, activities including protein synthesis and degradation occur segregated, far from the nucleus. Perhaps this functional segregation of signaling is a general mechanism by which cells conducting cellular activity in very fine structures far from the cell body. Further aspects of signaling will become more clear with the establishment of 3D cancer models which will offer more directions for imaging-based study of protrusion function [cite Bissell, ivaska] and the means by which we can effective block such function in the near future.

## Acknowledgements

We thank Darcie Babcock, Melissa Williams, and Dr. Claudia Lopez (FIB/SEM sample preparation and imaging; OHSU-FEI Living Lab), Aurelie Snyder (SIM, Advanced Light Microscopy Core, OHSU) for expert technical assistance, and Dr. Joe Dragavon (Biofrontiers Advanced Light Microscopy Core; CU Boulder; Howard Hughes Medical Institute) for N-STORM microscope use. The following funding sources made this work possible: ARCS Scholarship (Barbara and Phil Silver); W.Y.L.; NIH 5P50GM085273 and NIH/NCI P30CA118100, K.A.L.; NSF SBIR grants IIP-1353638, IIP-1644887, and IIP-1534745; Double Helix, SBIR IIP-1059286; K.H.; Damon Runyon Innovator Award and the Prospect Creek Foundation; X.N.; Oregon Nanotechnology and Microtechnology Institute NB3 Award, the Prospect Creek Foundation, and NIH RO1NS071116; T.Q.V.

## Supplementary Information

### Materials and Methods

#### Cell Culture

Human breast cancer cell lines SKBR3, MCF-7, and 21MT-1 were obtained from ATCC (Manassas, VA). Cell media and antibiotics were obtained from Gibco-Thermo Fisher (Waltham, MA). SKBR3 cells were grown in McCoy’s 5A containing 10% FBS. MCF-7 (low HER2-expressing, Luminal A) were grown in DMEM (10% FBS) and 21MT-1 (HER2 overexpressing) cells in RPMI (10% FBS) supplemented with 100 µg/mL streptomycin and 100 U/mL penicillin For live cell imaging or immunofluorescence labeling experiments, 100,000 cells were plated per 18 mm diameter, No. 1.5 glass coverslips (Electron Microscopy Sciences, Hatfield, PA) and cultured for 72 hours in 12-well plates (Corning Inc., Corning, NY) at 37°C in a humidified atmosphere of 5% CO_2_.

#### Live Cell Microscopy

SKBR3 cells were imaged in real-time on an inverted epifluorescence microscope (Zeiss Axiovert 200 M, 63x/1.4 oil objective, Oberkochen, Germany) equipped with an EM-CCD camera (iXon Ultra 897, Andor Technology, Belfast, Ireland) using Micro-Manager v1.4.17. Cells on coverslips were situated in a magnetic chamber (CM-B18-1, Quorum Technologies, Ontario, Canada) maintained at a constant temperature of 37°C using a heating inset (model P S1, PeCon GmbH, Erbach, Germany), a heating ring and temperature control unit (Temp Module S1, PeCon GmbH). For DIC microscopy, cells were imaged in serum-media (McCoy’s 5A with 10% FBS), and for HER2-QD imaging, cells were suspended in serum-media until immediately prior to imaging. During HER2-QD imaging, background and toxicity was minimized by imaging cells in FluoroBrite™ DMEM (Gibco-Life Technologies) buffered with 25 mM HEPES (Gibco-Life Technologies) and supplemented with 50 ng/mL ascorbic acid. DIC video microscopy was performed at a rate of 1 fps and fluorescence video microscopy was performed at rates of either 7 fps or 20 fps at a z-plane near the coverslip over 5000 frames. These dynamic motions were faster in time-scale compared to the motion of filopodial and lamellipodial extensions and retractions (**Movie 7**). An Optosplit II dual emission system (Cairn Research, Kent, UK) allowed simultaneous acquisition of two color channels (FITC and QD655) during fluorescence imaging.

#### Wide-Field Fluorescence Microscopy

For wide-field epifluorescence imaging, fixed SKBR3 cells were imaged using an inverted epifluorescence microscope (Zeiss Axio Observer.Z1, 63x/1.4 oil objective) equipped with an EM-CCD camera (Luca R 604, Andor Technology) using Micro-Manager. Cells were imaged over the total height of the cells by acquiring z-stacks at a z-step of 150 nm. Fluorescence emissions for Alexa488 and Alexa647 were detected using FITC and Cy5 filter cubes (excitation 480 nm/emission 535 nm and excitation 620 nm/emission 700 nm respectively; Chroma, Bellows Fall, VT), and QD fluorescence was detected using a QD655 filter cube (excitation 435 nm/emission 655 nm, Semrock, Rochester, NY). Image stacks were deconvolved using AutoQuant X2 (Media Cybernetics, Rockville, MD). For WGA-Alexa488, WGA-Alexa647, pAKT-Alexa488, and phalloidin-Alexa647, the 3D Blind Deconvolution algorithm was used with an adaptive PSF, medium noise value of 20, and 5 total iterations for 87 frames at a stack spacing of 150 nm. For HER2-QD655, the No/Nearest Neighbor algorithm was used with haze removal factor 0.99 and z-kernel width 2 for 87 frames at a stack spacing of 150 nm.

#### Structured Illumination Microscopy

For structured illumination microscopy (SIM), fixed SKBR3 cells were mounted in ProLong Gold antifade reagent (Life Technologies, Carlsbad, CA) and cured overnight at room temperature. Cells were imaged on a Zeiss Elyra PS.1 microscope with a 63x/1.4 oil objective equipped with an EM-CCD camera (iXon 897, Andor Technology) using ZEN software (Zeiss). Cells were imaged over a height of 3-5 µm from the coverslip by acquiring z-stacks at a z-step of 110 nm with 3 grating rotations. Fluorescence for WGA-Alexa488 was excited with a 488 nm laser, and emission was captured by a band-pass (BP) filter (495-575 nm) and long-pass (LP) filter (750 nm). Fluorescence for phalloidin-Alexa647 was excited with a 635 nm laser and emission was captured by an LP filter (655 nm). Images were reconstructed using ZEN to generate the final super-resolution SIM image.

#### 2D STORM Imaging

2D STORM imaging was performed on a custom PALM/STORM setup constructed on a Nikon Ti-U inverted microscope frame (Creech et al., 2017). Fresh imaging buffer containing an oxygen scavenging system and mercaptoethylamine (MEA) (Sigma-Aldrich, St. Louis, MO) was added 15 min before imaging. For 2D STORM imaging of Alexa647, both the 647 nm laser (1-2 kW/cm^2^) and the 405 nm laser (0-10 W/cm^2^) were applied to the sample simultaneously. The intensity of the 405 nm laser was increased gradually as the imaging progressed to ensure appropriate switching rates of the Alexa647 fluorophores. At these power densities, the EM-CCD was operated in frame transfer mode at 10 ms exposure time with an EM gain setting of 300, and each data set typically contained 50,000 frames. In all 2D STORM experiments, 100 nm gold nanoparticles were used as fiduciary markers. A custom-built focus stabilization system was used to maintain the image focus within ±25 nm. All image acquisition was done with Micro-Manager (Edelstein et al., 2014). Raw SMLM movies were processed using a custom-written MATLAB package (Nickerson et al., 2014).

#### Double-Helix 3D Super-Resolution Imaging

3D super-resolution imaging was performed on a Nikon N-STORM microscope with 100X 1.49 NA objective and the Double-Helix (DH) SPINDLE™ (Jain et al., 2016; Wang et al., 2017a) module with a Double-Helix phase mask (Grover et al., 2012; Pavani et al., 2009) optimized for far-red dyes. Images were captured using a Hamamatsu Orca Flash 4.0 sCMOS camera (2×2 binned) using a 30 ms exposure. The molecules were imaged using a 647 nm laser and the density of active molecules was maintained throughout the acquisition using a 405 nm laser to reactivate Alexa647 dye molecules. For each image 200,000 frames were acquired, resulting in more than 4 million localizations per reconstruction. Samples were imaged in freshly prepared buffer containing 1% beta-mercaptoethanol (\textbeta ME), 100 mM Tris, pH 8.0, 10 mM NaCl and an oxygen scavenging system (Dempsey et al., 2011). Individual fluorophores were localized in 3D using Double-Helix’s TRAX™ software. The data were drift corrected using the cross-correlation function and rendered using the normalized Gaussian method with the ImageJ plugin, ThunderSTORM (Ovesny et al., 2014). The z-depth was color coded using the ImageJ look up table (LUT) “Spectrum” (Schneider et al., 2012). The 3D super-resolution renderings of actin were created from tiff stacks of Control and Latrunculin B treated cells with 50 nm axial steps and 10 nm lateral steps that were imported into Imaris (Imaris X64 v 8.0.1, Bitplane Inc, Zurich, Switzerland). The video rotations (**Movie 17** and **Movie 18**) were created using the 3D view and animation functions in Imaris.

#### Volume Electron Microscopy Sample Preparation and Imaging

De-identified human breast tissue from a biopsy of a metastatic bone lesion was chemically fixed with EM-grade 2% paraformaldehyde and 1% glutaraldehyde in phosphate buffered saline (PBS). Separately, the bone lesion was characterized as HER2 positive by a pathologist based on hematoxylin and eosin stained slides of a parallel specimen 10% buffered formalin fixed and paraffin embedded. The specimen was processed for electron microscopy using a Pelco Biowave microwave (Ted Pella, Inc., Redding, CA) to assist with each step; all steps, unless indicated, were conducted at 150 watts, sometimes under vacuum. The specimen was first rinsed in buffer and then submerged in reduced 2% osmium tetroxide with 1.5% potassium ferricyanide at 100 watts for 3 min power ON, 2 min OFF, 3 min power ON, 2 min OFF, 3 min power ON. Specimen was rinsed 3 times in ddH_2_O and then submerged in aqueous 1% thiocarbohydrazide at 100 watts for 1 min power ON, 40 sec OFF, 1 min power ON. Following water rinses, the specimen was submerged in 2% osmium tetroxide at 100 watts for 1 min power ON, 40 sec OFF, 1 min power ON. Following rinses in ddH_2_O, the sample was submerged in 5% aq. uranyl acetate at 100 watts for 2 min power ON, 2 min OFF, 2 min power ON. The sample was then dehydrated through an ascending acetone gradient (50%, 75%, 95%, 100%, 100%) for 40 sec power ON at each step. The specimen was infiltrated with a 100% acetone and 100\% Epon resin mixture (1:1) for 3 min power ON and then with 100% resin four times for 3 min power ON before being polymerized in the oven at 60°C for 24 hours. Epon resin was made from Embed Kit (14120, Electron Microscopy Sciences, Hatfield, PA) mixture included: 73.5 g Embed 812, 45.5 g DDSA, 38.5 g NMA, and substituted 4.2 mL BDMA instead of DMP-30. 650 nm sections were cut, on a Leica Ultracut 7 (Leica Microsystems, Germany), and stained with toluidine blue to assess tissue architecture.

For FIB-SEM, resin-embedded samples were polished with a dry diamond knife tool to expose the area of interest on both the top and one side of the block and then mounted to a 45° pre-tilt SEM stub using colloidal silver paint. Blocks were sputter coated with 10 nm palladium with a Hummer. Milling and imaging of the block was performed on a FEI Helios Nanolab 660 Dual Beam FIB with AutoSlice and View software (FEI Inc., Hillsboro, OR). 6144 by 4096 pixel images were collected with the Elstar in-lens TLD detector in BSE mode at 2 kV with horizontal field width of 30 µm at a working distance of 3 mm; FIB milling was performed at 77 pA to generate a z-dimension step size of 5 nm – 1240 total slices for a complete depth of 6.2 µm. Volume representations, manual segmentations, measurements, and movie creations were performed using Amira software (FEI Inc., Hillsboro, OR).

#### HER2 Labeling using affiFAP-MG-QD for Live Single QD Imaging

HER2 was labeled by an affibody that binds the extracellular domain of HER2 (Z _HER2:342_, (Eigenbrot et al., 2010)) at the junction of domains III and IV. HER2 affibody (Z_HER2:342_) was fused to a fluorogen activating protein (FAP_dL5**_) at the N-terminus, C-terminus or both terminus (construct schematic and probe was prepared in the same manner as EGFR affiFAP, which were previously described in (Wang et al., 2015)). Three HER2 affiFAP constructs showed similar binding affinity to the fluorogen *in vitro* and to HER2 on SKBR3 cells. The HER2 affibody-FAP fusion (Z_HER2:342_-FAP_dL5**_), hereafter termed HER2 affiFAP, was selected for use in subsequent experiments. HER2 affiFAP was monovalently assembled to a quantum dot via selective binding of the FAP to its cognate fluorogen Malachite Green (MG) as previously detailed (Saurabh et al., 2014; Szent-Gyorgyi et al., 2008; Wang et al., 2017b; Wang et al., 2015). Biotinylated MG was pre-complexed with streptavidin-QD655 (Life Technologies) at a 1:1 stoichiometric ratio to a working concentration of 1 nM MG-QD655 for 15 mins at room temperature. HER2 affiFAP was added to live cells at 500 nM in 1% BSA and incubated for 10 mins in 37°C, 5% CO_2_. Following the HER2 affiFAP, pre-complexed MG-QD655 was diluted to 500 pM and labeled to cells for 5 mins at 37°C, 5% CO_2_. Immediately following labeling, cells were rinsed in PBS and mounted in imaging media of FluoroBrite™ DMEM (Gibco-Life Technologies) buffered with 25 mM HEPES (Gibco-Life Technologies) and supplemented with 50 ng/mL ascorbic acid to minimize photobleaching. A membrane marker (CellMask™ Plasma Membrane Green, Life Technologies) was added dropwise into the chamber containing cells at 0.03X prior to the start of live acquisition. Cells were maintained at a constant temperature of 37°C for the duration of imaging.

#### Immunolabeling of HER2, pAKT, and Actin in Fixed Cells

SKBR3 cells were fixed in 2% PFA containing 1 µg/mL of the membrane marker wheat germ agglutinin-Alexa488 (WGA-Alexa488, Life Technologies) or -Alexa647 (WGA-Alexa647, Life Technologies) for 10 mins at 37°C, then transferred to a fresh solution of 2% PFA for 20 mins at 37°C. For HER2 immunolabeling, cells were permeabilized in 0.1% Triton-X for 10 mins at room temperature, labeled with HER2 affiFAP (500 nM, 30 mins, 37°C) and incubated for 1 hr in blocking solution (8% NHS and 2% BSA at room temperature). MG-QD655 was pre-complexed as described above to a working concentration of 5 nM, incubated with free biotin at 50 nM for 5 mins at room temperature to occupy unbound streptavidin, diluted in blocking solution to 1 nM and added to cells for 1 hr at 37°C. Following, cells were rinsed in 2% blocking solution and mounted in borate buffer for image acquisition. To verify HER2-specific binding of the probe, cells were labeled following the same scheme with FAP alone (no affibody) and MG-QD655 as a negative control to ensure absence of non-specific binding. For pAKT immunolabeling, cells were permeabilized in methanol for 30 mins in three cycles (10 mins at −20°C, 10 mins at room temperature, 10 mins at −20°C), rehydrated in PBS for 20 mins at room temperature, and incubated for 1 hr in a blocking solution of 8% normal goat serum and 2% BSA. Cells were labeled with primary antibody to pAKT (Ser473, Cell Signaling Technology, Danvers, MA) diluted 1:100 in blocking solution (1.5 hr, 37°C), incubated for 1 hr in blocking solution, and then labeled with secondary antibody Alexa488 (goat anti-rabbit, Life Technologies) diluted 1:250 in blocking solution for 1 hr at 37°C. Following, cells were rinsed in 2% blocking solution and mounted in 1X PBS for image acquisition. To verify pAKT-specific binding, cells were labeled following the same scheme with no primary antibody and secondary antibody alone as a negative control to ensure absence of non-specific binding. For actin immunolabeling, cells were labeled with 500 nM phalloidin-Alexa647 (A22287, Life Technologies) overnight at 4°C and gently rinsed in PBS prior to mounting in PBS for image acquisition.

#### HER2 staining for 2D STORM

SKBR3 cells were cultured in 8-well µ-Slide chambers (ibidi, Fitchburg, WI) situated on No. 1.5 glass-bottom coverslips for 24-48 hrs at 37°C in a humidified atmosphere of 5% CO_2_ in McCoy’s 5A media supplemented with 10% FBS. Prior to plating, chambers were washed in 1 M NaOH for 2 hr at room temperature, rinsed with PBS five times, and incubated with PBS overnight. Cells were fixed in 3.7% PFA in 1X PHEM buffer for 20 mins, then rinsed with PBS. For HER2 immunolabeling, cells were blocked in 5% BSA for 30 mins, then labeled with 16 µg/ml Herceptin, a clinical monoclonal antibody that labels the extracellular domain of HER2, for 45 mins at room temperature. Following, cells were washed three times in PBS for 5 mins, then labeled with secondary antibody Alexa647 (goat anti-human, Life Technologies) diluted at 1:1000 (1 µg/ml) for 30 mins with protection from light. Cells were washed three times in PBS for 5 mins, and post-fixed with 3.7% PFA in 1X PHEM buffer for 10 mins. Following post-fixation, cells were washed with PBS and stored in the dark until imaging.

#### Actin staining for 2D SIM and 3D Super-Resolution

SKBR3 cells were cultured in MatTek dishes (MatTek, Ashland, MA) situated on No.1.5 glass-bottomed coverslips for 72 hrs at 37°C in a humidified atmosphere of 5% CO_2_ in McCoy’s 5A media supplemented with 10% FBS. Cells were prepared using a method developed by Xu et al. 2012. Briefly, cells were washed in cytoskeleton preserving buffer (CB buffer), then fixed and extracted in 2% glutaraldehyde and 0.25% Triton-X in CB buffer for 10 mins at 37°C. Cells were washed in CB buffer in 37°C, suspended in 0.1% NaBH_4_ in PBS for 15 mins to quench free glutaraldehyde, and washed in PBS two times. Tetraspeck™ Microspheres (0.1 µm, Invitrogen) were added to cells at a 1:200 dilution for 15 mins at 37°C to serve as fiducial markers, then cells were washed three times in PBS. F-Actin was labeled overnight at 4°C with Alexa647 conjugated Phalloidin (Invitrogen) at a concentration of 0.5 µM. Immediately before imaging, samples were rinsed once in PBS, then placed in imaging buffer (Xu et al., 2012).

#### Cell Treatment with NRG1-β1, Latrunculin B and G-Fascin

Cells were serum-starved overnight, then treated with 40 ng/mL neuregulin1-beta1 (396-HB/CF, R&D Systems Minneapolis, MN), 100 nM Latrunculin B (428020-1MG, Calbiochem-EMD Millipore, Billerica, MA), 125µM fascin-G2, a combination of 40 ng/mL neuregulin1-beta1 (NRG1) and 100 nM Latrunculin B (LatB), or a combination of 40ng/mL NRG1 and 125µM fascin-G2 for 15 mins at 37°C under a humidified 5% CO_2_ atmosphere. All treatments were diluted with serum-free media for application to cells. Cells were immediately fixed after treatment as described. For localized NRG1 application in live cell imaging, SKBR3 cells were serum-starved for 4 hr prior to NRG1 injection. Micropipettes were heat-pulled (PC-10 Dual-Stage Puller, Narishige, East Meadow, NY) from glass capillaries (outer diameter = 3 µm, 1B100F-4, World Precision Instruments, Sarasota, FL), gently positioned near cells (MHW-3 Manual Hydraulic Manipulator, Narishige) and vacuum injected (5 one-sec pulses at 5 kPa over 10 sec duration, IM-31 Microinjector, Narishige). Glass micropipettes were back-filled with 2.5 ng/µL NRG1 solution or 1X PBS and locally applied to a subcellular site of a cell during live acquisition. Control experiments using micropipettes filled with CellMask™ Plasma Membrane Green confirmed a localization of approximately 5-6 µm from the tip of the pipette immediately following vacuum injection (**Movie 4**). For Latrunculin B application during live cell imaging, cells were treated with 100 nM LatB during the affiFAP and MG-QD labeling steps (15 mins), then suspended in imaging solution containing 100 nM LatB in an imaging chamber for live acquisition. Cells were maintained at a constant temperature of 37°C for the duration of imaging.

#### HER2-siRNA Transfection

ells were transfected with HER2-targeting siRNA (Hs_ERBB2_14 FlexiTube siRNA, Qiagen, Hilden, Germany) or control non-silencing siRNA (AllStars Negative Control siRNA, Qiagen, Hilden, Germany) using siLentFect™ Lipid Reagent for RNAi (170-3360, Bio-Rad, Hercules, CA). siRNA and transfection reagent were diluted in 100 mM NaCl solution and incubated with cells plated at 50% confluency at 20 nM siRNA for 58 hr. Following siRNA transfection, cells were serum-starved for 6 hr, then treated with 40 ng/mL neuregulin1-beta1 for 15 mins, and fixed and labeled for HER2 as previously described.

#### Quantification of Filopodia-positive Cells Images

Filenames of images of siControl and siHER2 treated cells were scrambled, de-identified of their treatment conditions, and provided to a blind observer for assessment for the presence of filopodia in cells. Individual cells were graded at planes near or at the coverslip for the presence (Y) or absence (N) of 5 or more filopodia. Following, the percentage of cells containing 5 or more filopodia 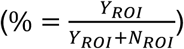 was measured for each ROI (n=60 ROI’s from 3 different experiments), linked back to its corresponding siControl or siHER2 group, and recorded for the prevalence of filopodia-rich cells in each condition.

#### HER2-QD and pAKT Quantification

For siRNA experiments, HER2-QD quantity per cell was measured for siControl or siHER2 treatment by quantifying the number of HER2-QD in the maximum projection of each image and dividing by the number of cells in the image 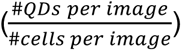. The HER2-QD count per image was quantified by custom MATLAB R2014B software (MathWorks), ‘QD Counter’, which created a maximum projection of a z-stack and located and counted the number of QDs present in the QD channel. The number of cells per image was determined by identifying the number of individual cells visible by WGA staining. For HER2-QD quantification in protrusions, images of WGA staining from cells on the plane of the coverslip were annotated in Fiji (Fiji is Just ImageJ) using the freehand selection tool to demarcate areas of protrusions or the whole cell at the same z-plane. Protrusions were identified on the basis of (1) their ruffled and discontinuous staining as evidence of cell membrane that contain extensive folds and elaborations, and (2) a greater than 1 µm distance between the extracellular edge of WGA stain and the cytoplasmic edge of the membrane label (in comparison to the approximately 1 µm distance observed in the continuous and “smooth” staining of membranes relatively low in protrusions). The entire cell was annotated by tracing the extracellular edge of WGA staining of each cell (entire cell denotes all membrane regions including protrusions and cytoplasmic regions). HER2-QDs were manually counted in the annotated regions using PointPicker in Fiji, where QDs displaying at least a 2×2 pixel size residing within the border and in contact with the annotated border was counted. HER2-QD surface density in protrusions and entire cell was calculated by dividing the sum number of QDs present in each annotated region and dividing by the region area 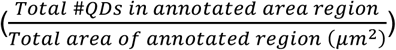. For pAKT-Alexa647 quantification, images of WGA staining from cells on the plane of the coverslip were annotated as previously described. pAKT-Alexa density in protrusions and entire cell were calculated by dividing the sum intensity of Alexa present in each annotated region and dividing by the region area 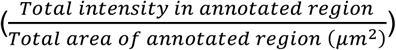. To compare the density of HER2 or pAKT in protrusions versus the entire cell, the density measurements from the two cellular locations were compared by a protrusion to entire cell ratio. The protrusion-to-entire cell ratio was calculated by dividing the density of HER2 in the protrusion by the density of HER2 in the entire cell for each individual cell. 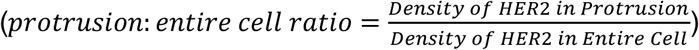 The protrusion-to-entire cell ratio for pAKT was similarly calculated.

#### Western Blot

Cells were lysed in a RIPA buffer (Sigma-Aldrich) with Halt Protease and Phosphatase Inhibitor Cocktail (Pierce-Thermo Scientific) and then cleared by centrifugation. Protein concentration was estimated with a BCA assay (Pierce-Thermo Scientific). Proteins present in cell lysates (50 µg) were resolved by SDS-PAGE and transferred onto Immobilon-FL PVDF membrane (EMD Millipore). Membranes were probed with antibodies specific for HER2 (3B5, Sigma-Aldrich) and Actin (sc-1615, Santa Cruz Biotechnology, Dallas, TX). Immunoreactive proteins were detected and quantified using infrared fluorescent IRDye® secondary antibodies and Odyssey® imagers (LI-COR Biosciences, Lincoln, NE).

#### Single particle QD Tracking

Single particle tracking of quantum dot-tagged HER2 receptor complexes was performed as previously described (Valley et al., 2015; Vermehren-Schmaedick et al., 2014). Briefly, we applied custom-written MATLAB algorithms to subtracting the camera offset and dividing by a gain factor to convert image data from raw output to Poisson distributed ‘counts’ as previously described (Lidke et al., 2005b; Smith et al., 2010). Areas in each image were identified as possible candidates for single particle fitting using a difference of Gaussian filtering step as described in (Huang et al., 2011) and each candidate area was fit using the maximum likelihood method described in (Smith et al., 2010). Fits that were above a threshold intensity and matched the expected point spread function shape (log-likelihood ratio test described in (Huang et al., 2011)) were connected into trajectories. Trajectory connection was performed using a modification of the cost matrix approach (Jaqaman et al., 2008). Trajectories were visually inspected to verify absence of connection errors before further analyses. HER2 trajectories were assigned to filopodia, lamellipodia, and cell body based upon the location of the receptor track relative to the membrane label. Maximum intensity time projections of the HER2-QD channel was used to identify filopodia that were difficult to resolve by membrane label alone. Trajectories corresponding to slender filaments less than 900 nm in diameter and more than 2 µm in length (as visualized by HER2-QD time projection) from the edge of the membrane label were classified as filopodia. Trajectories along characteristically broad membrane sheets that extended at least 3 µm from the cell body were classified as lamellipodia, and trajectories along regions of the membrane appose to the bulk cytoplasm were classified as cell body. HER2-QD was also found in motion in cell regions that could be mistakenly interpreted as cytoplasmic but were on the ‘underside’ of cells at the plane of the cover slip. This was confirmed by: 1) real-time focusing of dynamic HER2-QDs showing that they are in the same plane of focus as single blinking QDs adherent to the coverslip and 2) 3D deconvolution of cells fixed following conditions similar to real-time tracking that showed the presence of HER2-QDs at the plasma membrane at the plane of the coverslip and absence of HER-QDs in the cell cytosol. To avoid potential artifacts introduced by interference of the motion of HER2-QDs with the coverslip, these trajectories were not included in our analysis.

#### Statistical Analysis of HER2-QD Motion

All analyses were performed in MATLAB (Mathworks, v. 7.10.0, Natick, MA). <u>*1D analysis of filopodial motion:*</u> 2D filopodia data were transformed by rotation to minimize their root mean square deviation along the y-coordinate. Subsequently, the x-coordinate was used for analysis. *<u>Simulations:</u>* Simulations of Brownian dynamics were performed in one and in two dimensions. Step sizes were successively drawn from a 1D and a 2D normal distribution, respectively. Simulation lengths (1D: 10 trajectories of 820 steps; 2D: 23 trajectories of 1000 steps) and the normal distribution variances (1D: σ^2^=0.0625; 2D: σ^2^=0.005) correspond to the trajectory lengths and the average variance as derived from experimental mean-square displacement data. In the 1D simulations, the particle is confined to the region [-2,+2] to reflect the representative average length of a filopodia (∼4 μm). Simulation of 2D random walks were unbounded. <u>*MSD, autocorrelation, step angle, and instantaneous velocity analysis:*</u> Mean square displacements versus time intervals were defined as <Δx(τ)^2^> = <(x(t+τ) - x(t))^2^> with lag time τ calculated over all jumps and all trajectories. Average values are calculated over all data points in all trajectories. A 1000-fold bootstrap sampling procedure was performed on each set of trajectories and 90% confidence intervals are reported as error bars at any data point(Efron and Tibshirani, 1994). The effective diffusion constant of each trajectory was determined using maximum likelihood estimation (D_MLE_) that takes into account the experimental effects of finite exposure time of the camera, varying localization precision and intermittent trajectories (Relich et al., 2016). We used auto-correlation to determine if correlated active transport was present in trajectories (Zuckerman, 2010). The first-step auto-correlation is defined as <Δx(t) * Δx(t+τ)> / <Δx^2^>. The first-step angle was measured as the angle between two neighboring steps. Instantaneous speeds of trajectories were computed as the mean single-jump velocity within a sliding window containing six localizations. In all experiments, the QD location at certain times may be unknown because of blinking, which results in some trajectories with missing coordinates at those time points. For the analysis of step sizes, angles, and auto-correlations, only those steps were considered that had coordinates recorded at a constant time interval τ = 0.13 s. Except for the step angle, which is not defined in 1D, all analysis graphs are shown in 1D (i.e. the average of the x and y components of 2D data) for a correct comparison to the filopodia and the 1D simulation data.

#### Other Statistical Analyses

Data in each experiment was evaluated by ‘qqplot’ function in MATLAB R2014b to test for the normality of data. For data in which the normality assumption was not met, the non-parametric Mann-Whitney rank sum test was conducted (‘ranksum’ function in MATLAB) to test the null hypothesis that data from two groups come from continuous distributions with equal medians against the alternative that they do not. For multiple comparisons, the Mann-Whitney rank sum test was conducted between relevant groups with a Bonferroni adjustment of α based on the number of comparisons at a p<0.05 significance cut off. The Bonferroni adjusted p-value was determined by dividing 0.05 by the number of comparisons. For comparing the equality of cumulative probability distributions of HER2 dynamics in different regions, distributions of diffusion constant, and instantaneous velocity, the non-parametric Kolmogorov-Smirnov test was conducted (‘kstest2’ in MATLAB) to test the null hypothesis that data from two groups come from the same continuous distribution at a p<0.05 significance level. For comparing the linear correlation between HER2 expression and percent of filopodia positive cells, Pearson’s correlation test was conducted (‘corr’ in MATLAB) to test the null hypothesis of no correlation between the two variables at a p<0.05 significance level and for computing the Pearson’s correlation coefficient.

**Figure S1:**
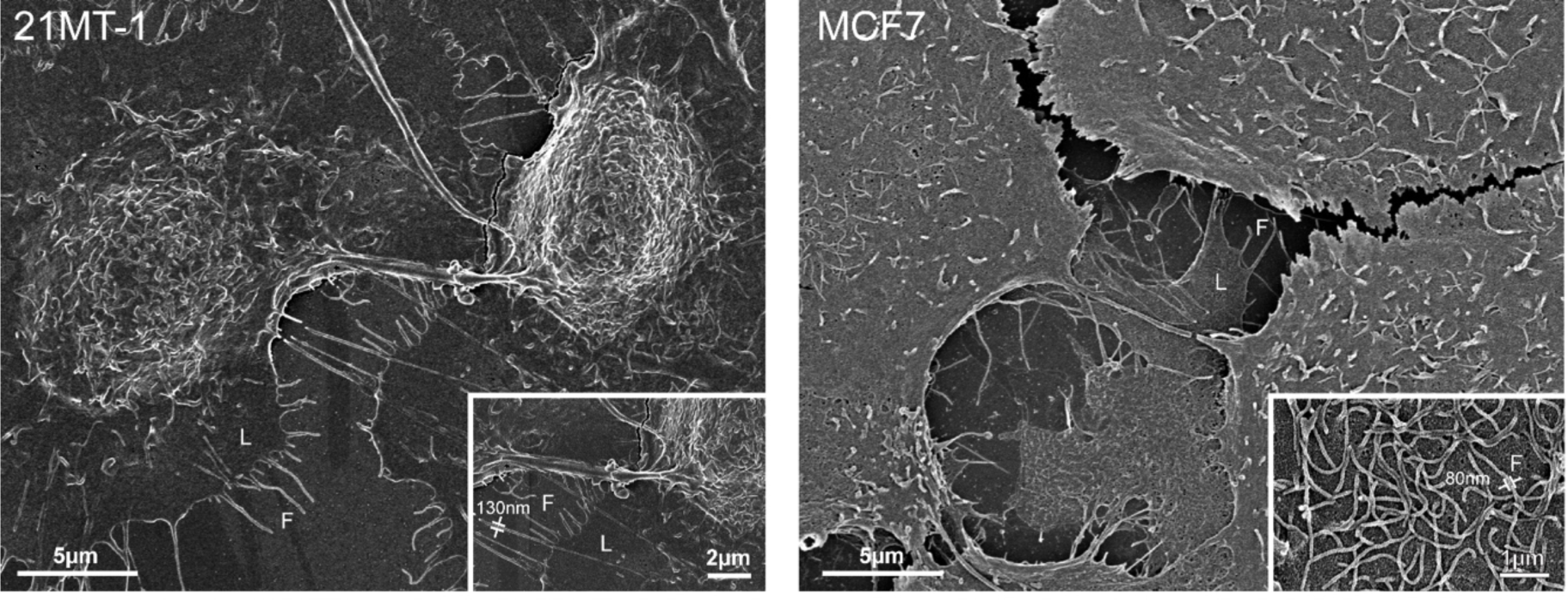
Segregated, far-reaching morphology of filopodia and lamellipodia are a common feature of additional human breast cancer epithelial and luminal cell models. SEM of slender filopodia (F) extending laterally from lamellipodial sheets (L) in high-HER2 expressing 21MT-1 breast cancer epithelial cell (*left*) and low HER2-expressing MCF7 (*right*) human breast cancer luminal cell models. Cultured in serum-media, *in vitro*.

**Figure S2:**
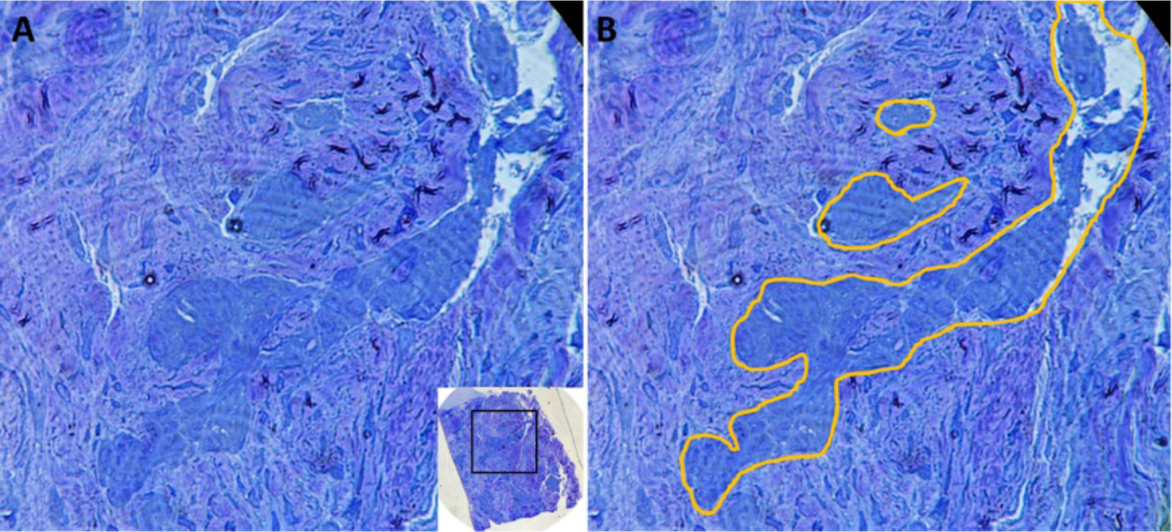
Toluidine-blue stained 600 nm thick resin section of the *in vivo* HER2-positive breast tumor cell obtained from a biopsy of a metastatic bone lesion that was prepared for Volume EM. **(A)** The mass of breast cancer cells is observed infiltrating old bone. Inset is low magnification of whole section. **(B)** The breast cancer infiltrate is outlined in orange.

### SI3 Additional information about HER2-QD probe characterization

We verified the functionality of various versions of HER2 affibody-FAP fusion binding to malachite green (MG) and characterized them according to methods previously described for the EGFR affibody-FAP fusion (Wang *et al.*, 2015). The FAP was fused to the C-terminus (Z_HER2:342_-FAP_dL5_**, AF) or N-terminus (FAP_dL5_**-Z_HER2:342_, FA) of the affibody, or between the dimeric affibody (Z_HER2:342_-FAP_dL5_**-Z_HER2:342_, AFA). Monomeric proteins of affibody (Z_HER2:342_, A) and FAP (FAP_dL5_**,5) alone were used as controls (**Figure S3A**). The fluorescence spectral properties of the HER2 affibody-FAP fusion to MG and the various constructs were analyzed. FAPs have the capability to rapidly associate with non-fluorescent MG for thousands-fold fluorescence enhancement (Szent-Gyorgyi *et al.*, 2008). In the presence of MG-Btau, the affibody-FAP fusions: AF, FA, and AFA all displayed similar fluorescence spectra to F alone, indicating fusion of affibody did not alter FAP activation of MG fluorescence enhancement (**Figure S3B**). We compared the fluorescence activation of MG by HER2 affibody and FAP; only MG binding to FAP yielded fluorescence enhancement, while MG binding to affibody alone did not (**Figure S3C**). We next confirmed that FAP and MG association was unaffected by the fusion of the HER2 affibody. Binding equilibrium analysis of recombinant probes demonstrated that HER2 affibody-FAP fusions and MG have similar association rates compared to F and MG alone. These fluorescence measurements under equilibrium conditions indicate that the three affibody-FAP fusion probes have nanomolar dissociation constants that are comparable to F alone (**Figure S3D**). Thus, these data show that the fluorogen activating properties of the affibody fusion FAP is maintained and unchanged from the unmodified FAP.

We next validated that the conjugation of FAP preserves HER2 affibody binding specificity for HER2. Dissociation constant analysis of HER2 affibody-FAP on SKBR3 cell surface show high affinity binding of HER2 affibody-FAP to HER2. All probes demonstrated low KD measurements, indicating high affinity binding of the affibody-FAP fusion to HER2 (**Figure S3E**). FAP conjugation to the HER2 affibody did not affect the binding specificity to HER2 as shown by a competition assay between nonfluorescent HER2 affibody (A) and the HER2 affibody-FAP fusion protein (AFA) for HER2 binding (**Figure S3F**). HER2 affibody-FAP fusion detected by MG was confirmed to label HER2 in SKBR3 (**Figure S3G**). These data show that the HER2 affibody fusion preserves the binding specificity of the affibody for HER2.

Finally, we utilized the HER2 affibody-FAP-MG system in conjunction with a QD for monovalent and specific labeling of HER2 with single QDs in SKBR3. First, HER2 affibody-FAP fusion, AF (Z_HER2:342_-FAP_dL5**_), was added to cells for labeling of HER2 (step 1). Second, to enable single-molecule imaging of HER2, HER2 affibody-FAP (HER2 affiFAP) was visualized using a streptavidin QD655 and biotinylated MG (step 2) conjugated at 1:1 ratio (**Figure S3H**). The FAP-MG system allows the capability to tune the number of receptors detected by adjusting the quantity of MG-QD added. MG-QD655 conjugate was added at a 500-fold lower concentration than the affiFAP to ensure single QD detection of HER2. HER2-QDs were a composition of dimerized and single HER2 receptors. For simplicity, HER2-QD will refer to HER2 labeled by HER2 affibody-FAP-MG-QD unless otherwise specified. Single QD detection of HER2 was enabled by the discrete and punctate QD signal and was confirmed by representative fluorescent blinking profiles (yellow plots) from two HER2-QDs which show the characteristic on-off behavior typical of single QDs. Time lapse imaging was conducted of stationary HER2-QDs in fixed cells at a single plane on the coverslip for 50 sec at 20 frames per second. Profiles of total intensity of individual QD puncta (2×2 pixels) were plotted over time (n = 21 QDs analyzed). These blinking profiles confirm that HER2 receptors were detected as single QDs (**Figure S3I**). HER2-QD was validated for HER2 specificity by comparing HER2-QD in HER2-positive SKBR3 and HER2-negative MCF7 and MDA-MB231. Fixed cells were labeled with 500 nM HER2 affiFAP, followed by 1 nM MG-QD655 as described in **Methods**. To verify HER2-specific binding of the probe, 500 nM FAP (no HER2 affibody), followed by 1 nM MG-QD655 was labeled to cells as negative control to ensure absence of non-specific binding (**Figure S3J**). The number of HER2-QDs per cell was quantified based on the maximum projection of the QD channel for each cell within each image. Altogether, these data show that our characterized HER2-QD probe has been validated to bind with monovalency and specificity to HER2.

**Figure S3:**
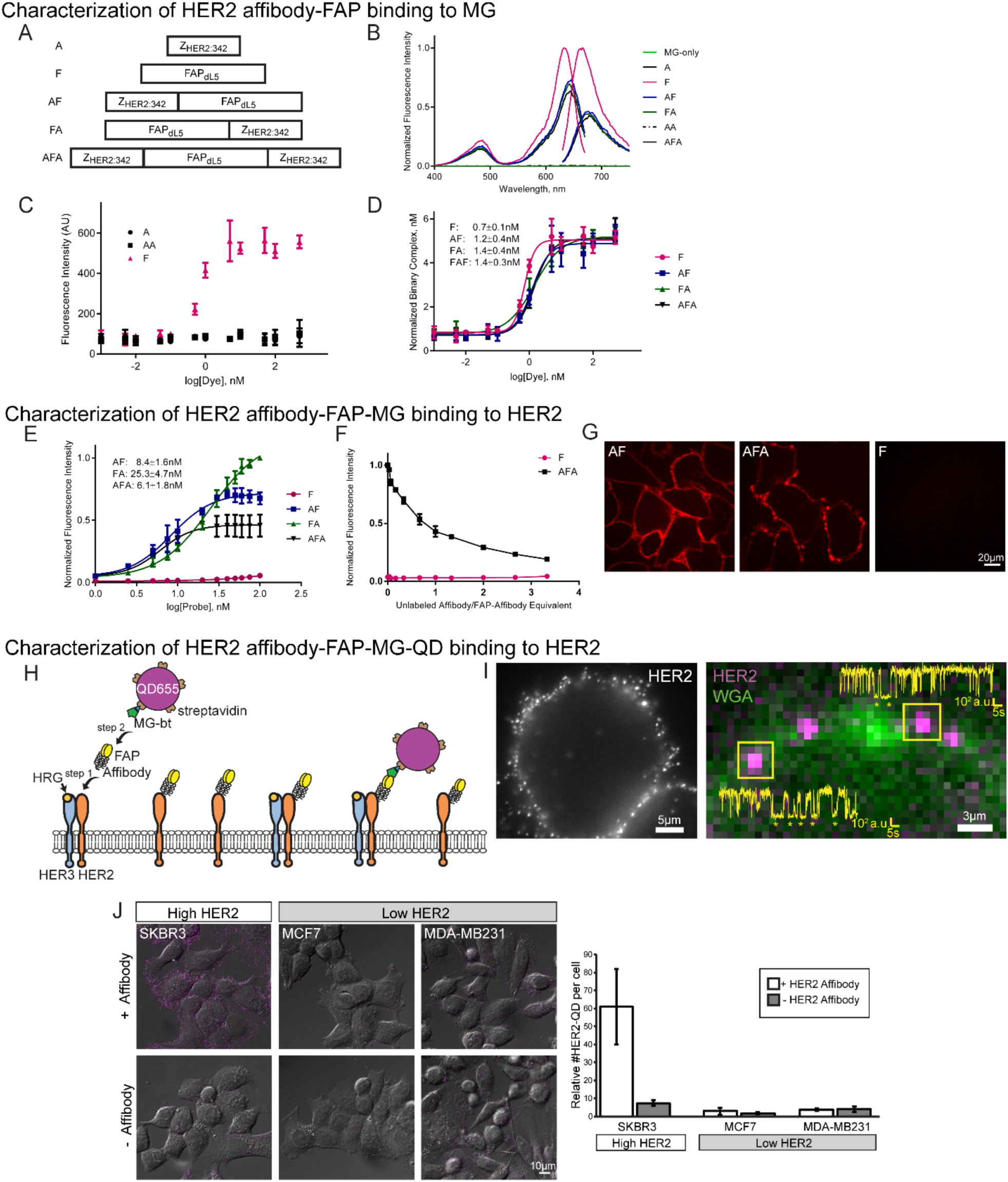
Characterization of HER2 affibody-FAP-MG-QD probe for monovalent and specific HER2 label. Affibody-FAP fusion constructs. **(B)** HER2 affibody-FAP fusion and MG functionality confirmed by fluorescence spectra. MG-Btau (1 µM), cell-impermeant analog of MG, was pre-complexed to probe constructs (10 µM). **(C)** MG binding to FAP yields fluorescence enhancement, while MG binding to affibody alone did not. MG-Btau titrated with 5nM of affibody Z_HER2:342_ (A), FAP_dL5**_ (F), or dimeric affibody (AA). Fluorescence measured at 636 nm (excitation) and 664 nm (emission). **(D)** FAP and MG association is unperturbed by HER2 affibody fusion to FAP as shown by binding equilibrium analysis. Fluorescence intensity corrected for fluorogen-only background, then normalized to FAP-MG fluorescence (1 nM). **(E)** HER2 affibody-FAP binds with high affinity to HER2 as shown by dissociation constant analysis of HER2 affibody-FAP on SKBR3. Mean fluorescence was corrected with FAP-MG background in cells, then normalized to mean fluorescence of probes (250 nM). **(F)** FAP conjugation preserves specificity of HER2 affibody to HER2 as shown by competition assay of nonfluorescent HER2 affibody (A) binding to cell surface. Cells labeled with AFA or F (250 nM), followed by serial dilution of A. MG-Btau (100 nM) added prior to measurement. **(G)** HER2 affibody-FAP fusion probes and MG display HER2 labeling in live SKBR3 cells, while FAP alone did not. HER2 detected by MG in cells labeled with fusion probes (30 nM) and MG-Btau (40 nM). **(H)** Two-step monovalent labeling of HER2 with single QDs in SKBR3. HER2 affibody-FAP fusion, AF (Z_HER2:342_-FAP_dL5**_), was added to cells (step 1) and detected by streptavidin QD655 and biotinylated MG (step 2) conjugated at 1:1. **(I)** Discrete HER2-QD fluorescence enables single QD detection of HER2. HER2-QD in fixed SKBR3 imaged at the plane of the coverslip (*left*), and blinking profiles (yellow, on *right*) of two HER2-QDs (magenta) on WGA-Alexa488 membrane (green) show characteristic on-off behavior of single HER2-QD probes. *y*-axis: intensity, *x*-axis: time, * are QD at off-state. **(J)** Validation of HER2-QD specificity. Max projection of HER2-QD on DIC of HER2-positive SKBR3 cells, and HER2-negative MCF7 and MDA-MB231 cells (*left*). HER2-QD quantity per cell plotted as mean ± SD (*right*, n=30 cells).

### SI4 Additional HER2 localization experiments conducted over longer durations following NRG1 stimulation

Fluorescence images were quantified to determine HER2-QD localization in membrane or cytoplasm following increasing durations of NRG1 stimulation. To understand whether HER2 undergoes internalization following receptor stimulation, we designed a protocol to label live cells with HER2 affiFAP prior to stimulation and to tag MG-QD only after fixation and permeabilization to (1) avoid labeling-induced internalization that may result following long stimulation durations, and (2) to identify HER2 receptors that remained membrane bound or HER2 receptors that internalized. SKBR3 cells were serum-starved overnight, labeled with HER2 affiFAP (500 nM for 10 mins at 37°C), then stimulated with HRG for 5 mins, 15 mins, or 1 hr. Following, cells were fixed and permeabilized, then labeled with MG-QD (as described for fixed cells, **Methods**). The number of HER2 in the membrane and cytoplasm were quantified for the mid-volume optical sections (across a depth of 5-7 µm depending on the height of cells) of each cell. The percentage of HER2 receptors residing on the membrane were consistently above 90% even with increasing lengths of stimulation, suggesting HER2 undergoes little if any internalization. Consistent with the percentage, HER2 receptor count on the membrane remains very similar even with varying lengths of stimulation, and the receptor count in the cytoplasm remains consistently small, indicating that HER2 remains primarily membrane bound and does not undergo internalization. The consistency in receptor count across different time points also confirms that the same number of receptors were retained over the duration of stimulation, and that no receptors were lost or gained. We confirmed HER2 receptor location by a secondary method using the fluorescence of the affiFAP-MG complex without the QD. SKBR3 cells were labeled with HER2 affiFAP, then placed in serum-containing conditions for 1 hr. Following, cells were fixed and permeabilized, then labeled with a membrane impermeant MG (MG-Btau). As observed with HER2-QD, HER2-MG also resided on the membrane and displayed little internalization. Altogether, our studies show that recycling and degradation of HER2 in the cytoplasm is uncommon.

**Figure S4:**
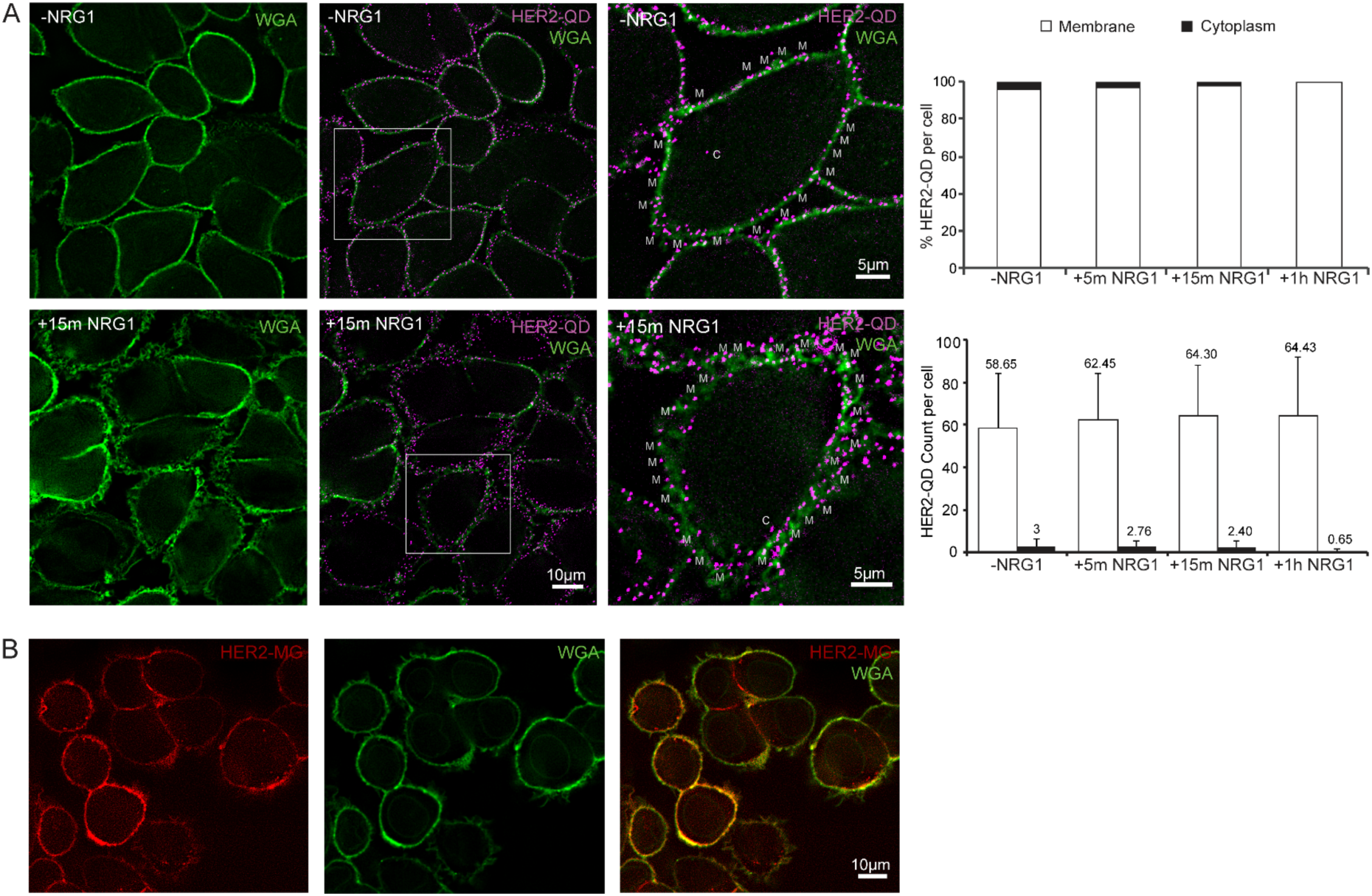
Activated HER2 receptors localize to membrane protrusions and remain primarily membrane bound. **(A)** HER2 receptors localize to the membrane and undergo little internalization. HER2-QD (magenta) and WGA-Alexa488 membrane (green) of untreated (-NRG1, *top row*) and 15 mins NRG1 stimulated (+15m NRG1, *bottom row*) SKBR3 cells at the mid-section of cells. Magnified view of cells from -NRG1 and +15 min NRG1 (third column) show that most HER2-QDs align on membrane label (M) and few receptors reside in the cytoplasm (C). HER2 quantity in membrane and cytoplasm for the mid-volume optical sections (across a depth of 5-7 µm) of each cell is plotted as mean percentage of HER2 per cell in each region (bar plot on *top right*) or as mean count of HER2 per cell in each region (bar plot on lower right). Error bars: SD, n=68-70 cells for each condition. **(B)** HER2 residence on membrane is additionally confirmed by HER2-MG fluorescence. HER2-MG (red) and WGA-Alexa488 membrane (green) in SKBR3 cells in serum-containing media for 1 hr. Similar to results from QD-detection of HER2, MG-detection confirmed HER2 receptors reside in large part on the membrane, and that there is little internalization of the receptor. Our studies show that recycling and degradation of HER2 in the cytoplasm is a rare event.

**Figure S5:**
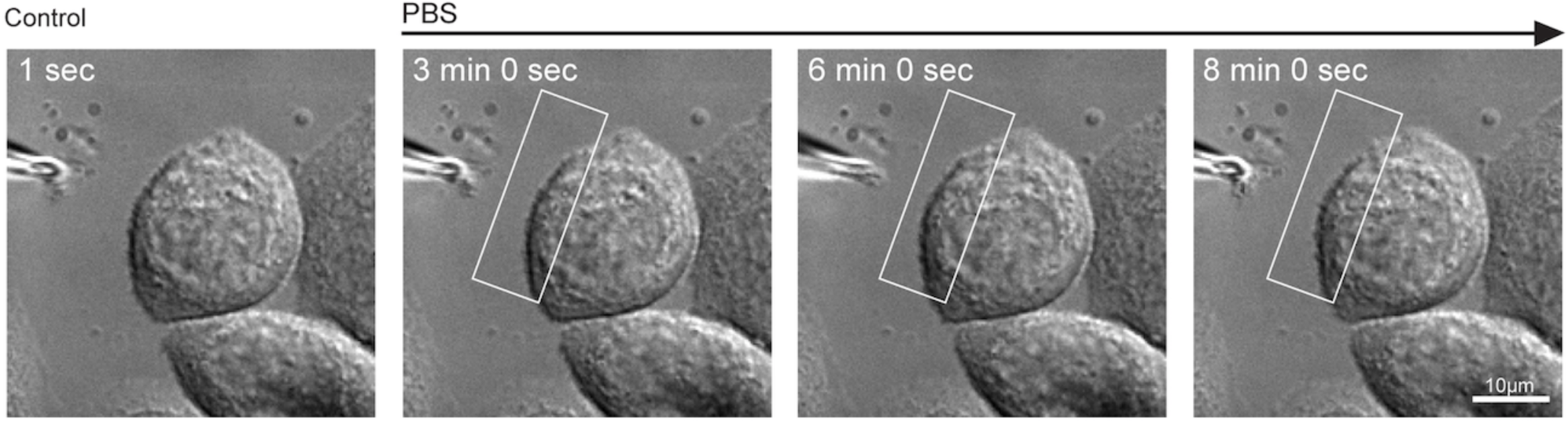
Control experiment for localized pipette delivery. Localized micropipette delivery of NRG1-free PBS does not evoke protrusion formation.

**Figure S6:**
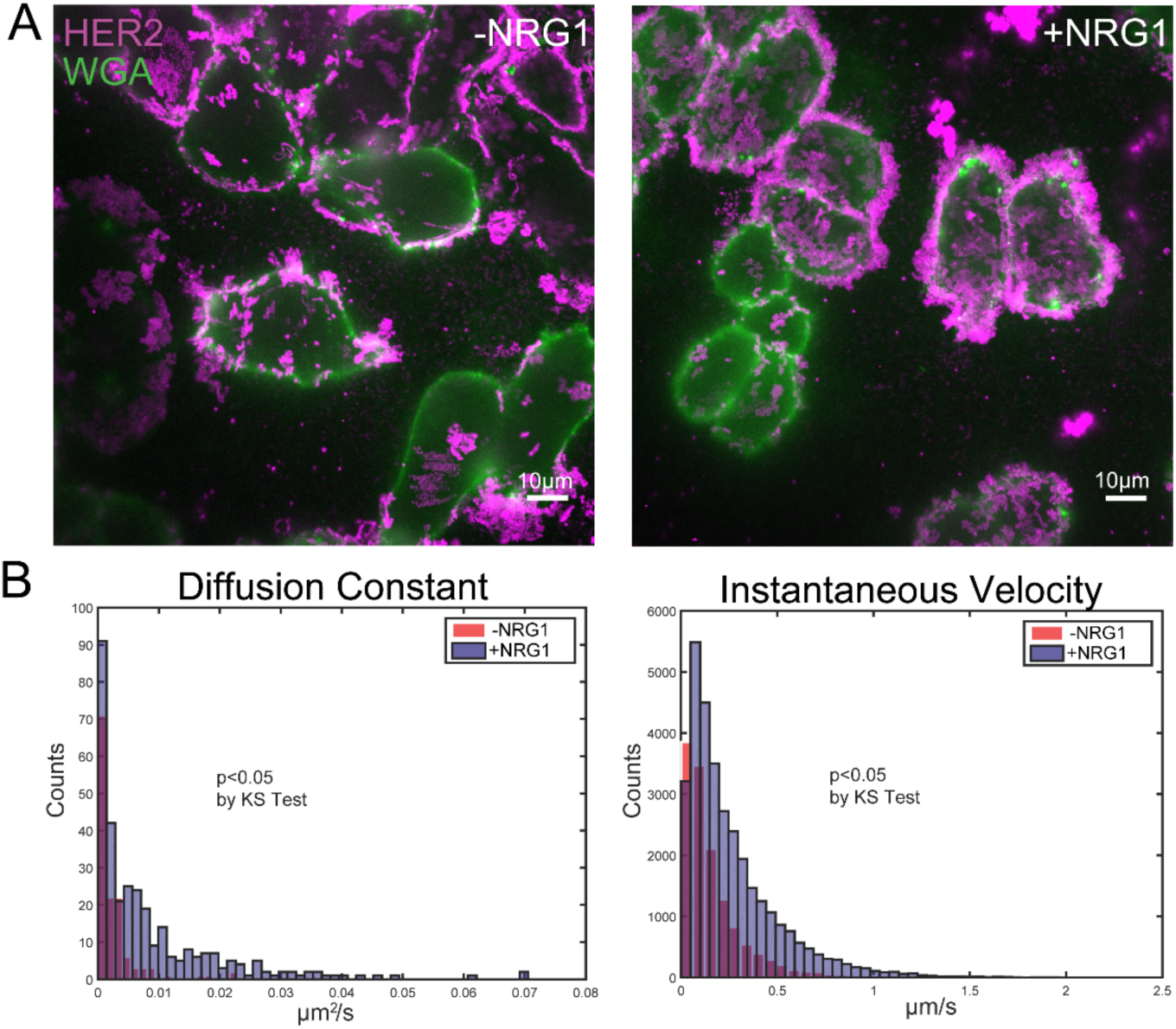
HER2 dynamics increase in mobility and instantaneous velocity upon NRG1 stimulation. **(A)** Untreated (-NRG1) and NRG1 stimulated SKBR3 cells (+NRG1) demonstrate qualitative differences in dynamics. Maximum projection of HER2-QD (magenta) fluorescence over time overlaid on still-image of membrane (green, acquired at start of acquisition). **(B)** Histogram of diffusion constant and instantaneous velocity for HER2 receptors in -NRG1 and +NRG1 stimulated cells. HER2 receptors in +NRG1 stimulated cells exhibit higher diffusion constants and faster instantaneous velocity. Counts are values of the diffusion constant and velocities computed from trajectories tracked in protrusion and non-protrusion regions of cells.

**Figure S7:**
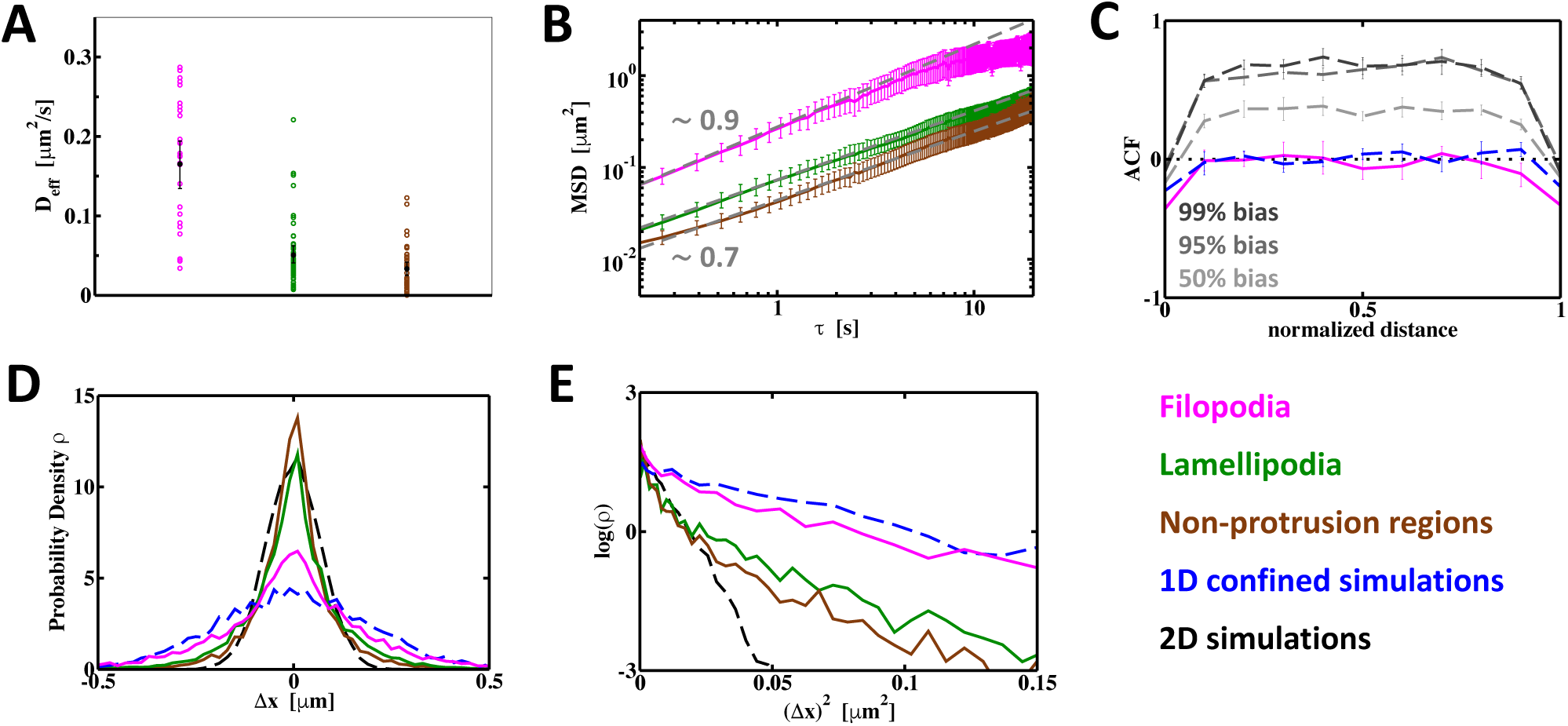
HER2 motion along the longitudinal axis of filopodia is diffusive Brownian motion. **(A)** The effective diffusion coefficient (D_eff_) of HER2 in filopodia (magenta), lamellipodia (green), and non-protrusion regions (brown) derived from linear fits to the initial data points of the corresponding MSDs (Figure 5A) as previously described (Michalet, Phys. Rev. E 83, 059904, 2011; Ernst and Kohler, Phys Chem Chem Physics, 2013). **(B)** Log-log plot of the HER2-QD MSDs as a function of the time interval τ (Figure 5A). The dashed grey lines show the slope of the individual MSDs (∼0.9 for filopodia, ∼0.7 for lamellipodia and non-protrusion regions) at short τ. **(C)** Two-step ACFs of as a function of the normalized distance traversed in confined 1D simulations (*dashed blue*). As a proof of concept for the use of these ACFs to identify a preferred direction, we performed the same 1D simulations with a bias that would keep the direction of particle movement to different degrees (50%, 95%, or 99% of the time). The corresponding ACFs show positive correlation to different degrees. Such a result can be expected from uni-directional and active motor-driven transport and is in stark contrast to the filopodial ACFs (magenta). **(D)** The distribution of HER2 step sizes, Δx, in the different cell regions and simulations indicates non-Gaussian behavior, i.e. non-Brownian motion, of HER2 in the lamellipodia and the non-protrusion regions. **(E)** The corresponding linearized distribution of step sizes, i.e., log(ρ) versus (Δx)^2^, reveals the deviation from linearity, and thus non-Gaussian behavior of HER2, in the lamellipodia and the non-protrusion regions, particularly when comparing to the 2D simulation.

**Figure S8:**
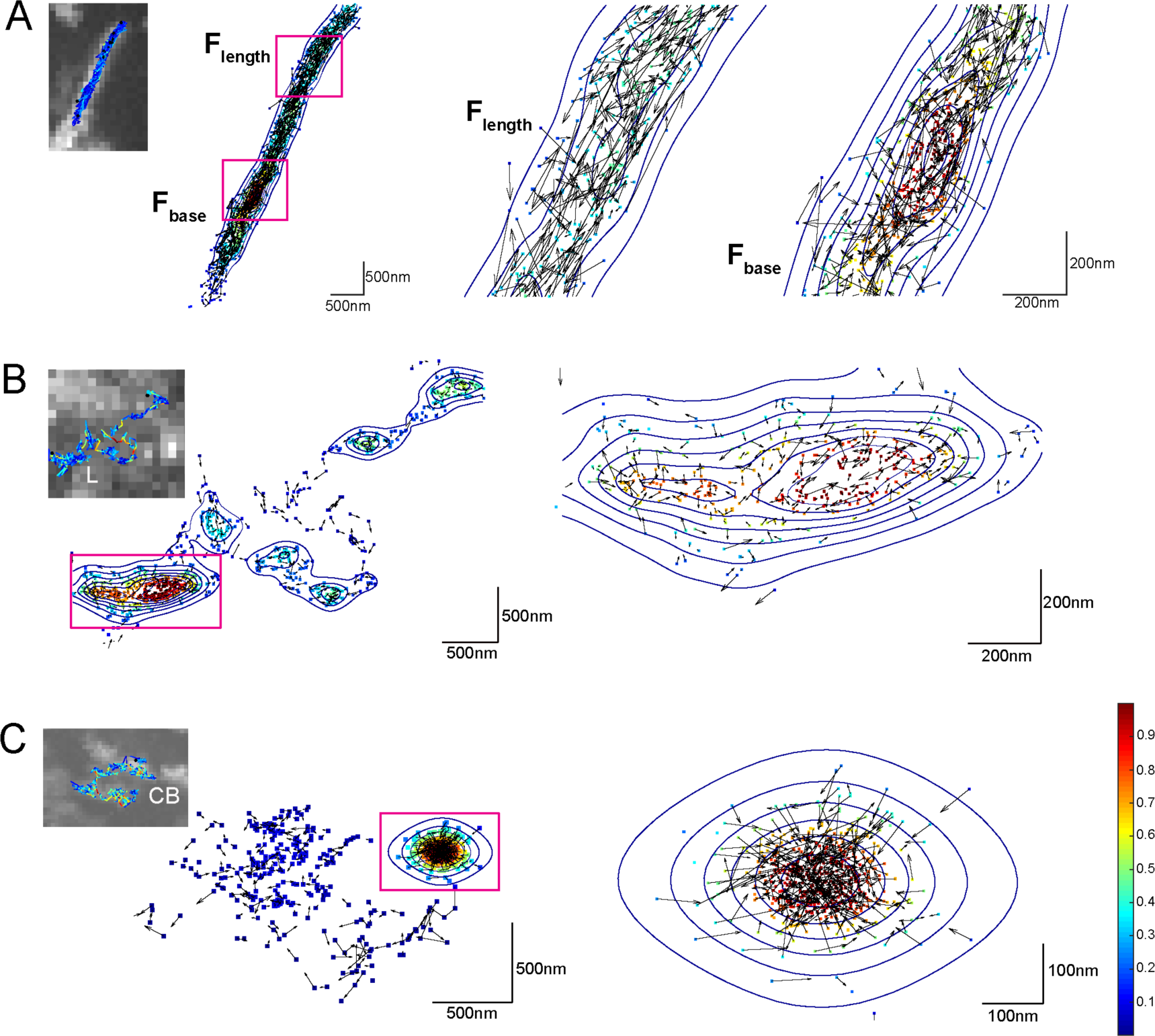
Additional examples of unimpeded HER2 diffusion along filopodial and confined HER2 diffusion in lamellipodia and the cell body. Maps showing the density contour of HER2 motion along with speed and direction attributes. Density contour is color-coded and normalized to each trajectory and velocity vectors are denoted by arrows with component (v_x_, v_y_) at the points (x, y). These maps reveal nanoscale membrane compartmented motions and directly show reflected motion of HER2-QDs at these compartment boundaries at the base of filopodia (A), lamellipodia (B) and the cell body (C).

**SI9.**
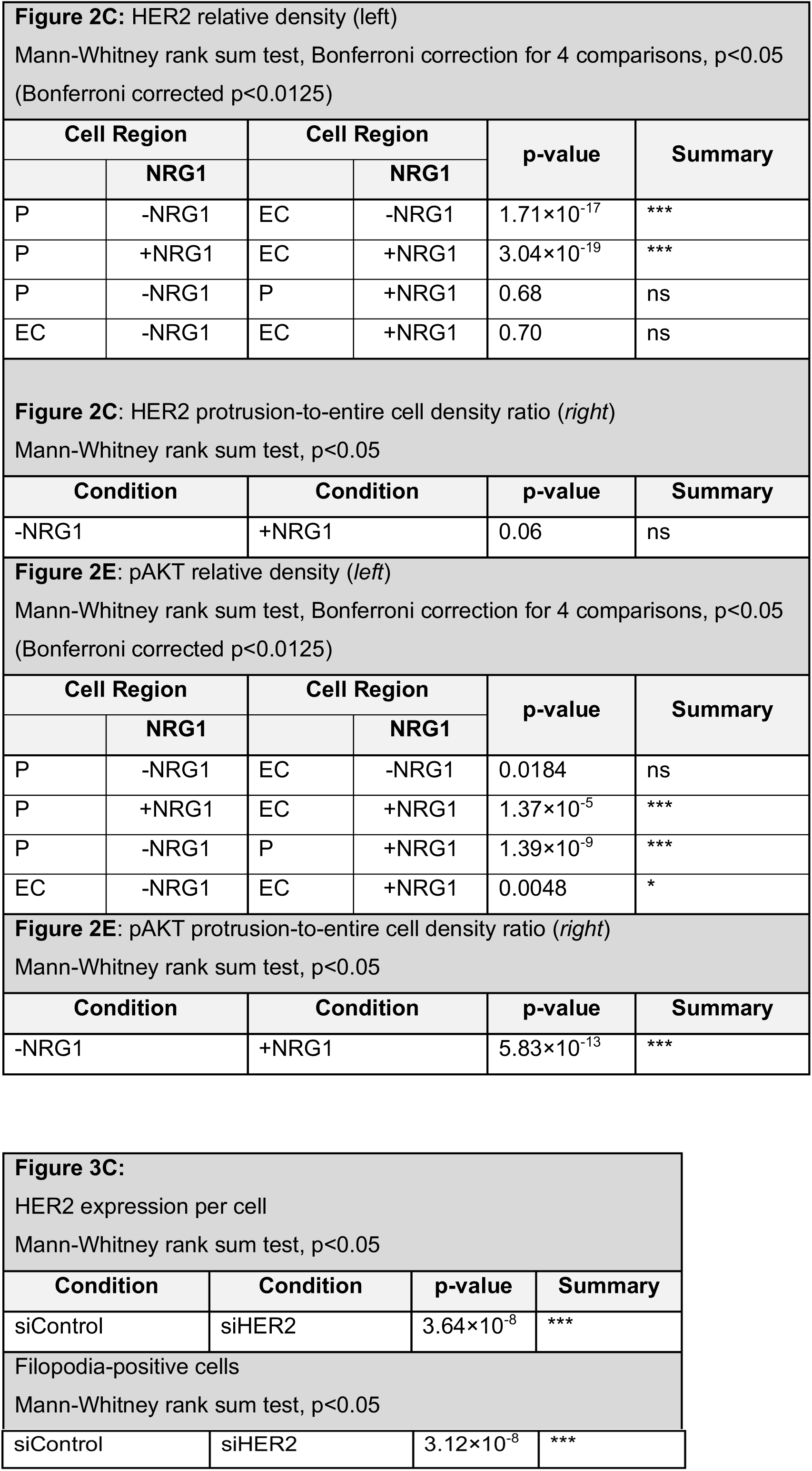
Test of significance for HER2 expression and filopodial growth; HER2 and pAKT density in cellular regions.

**STable 1.**
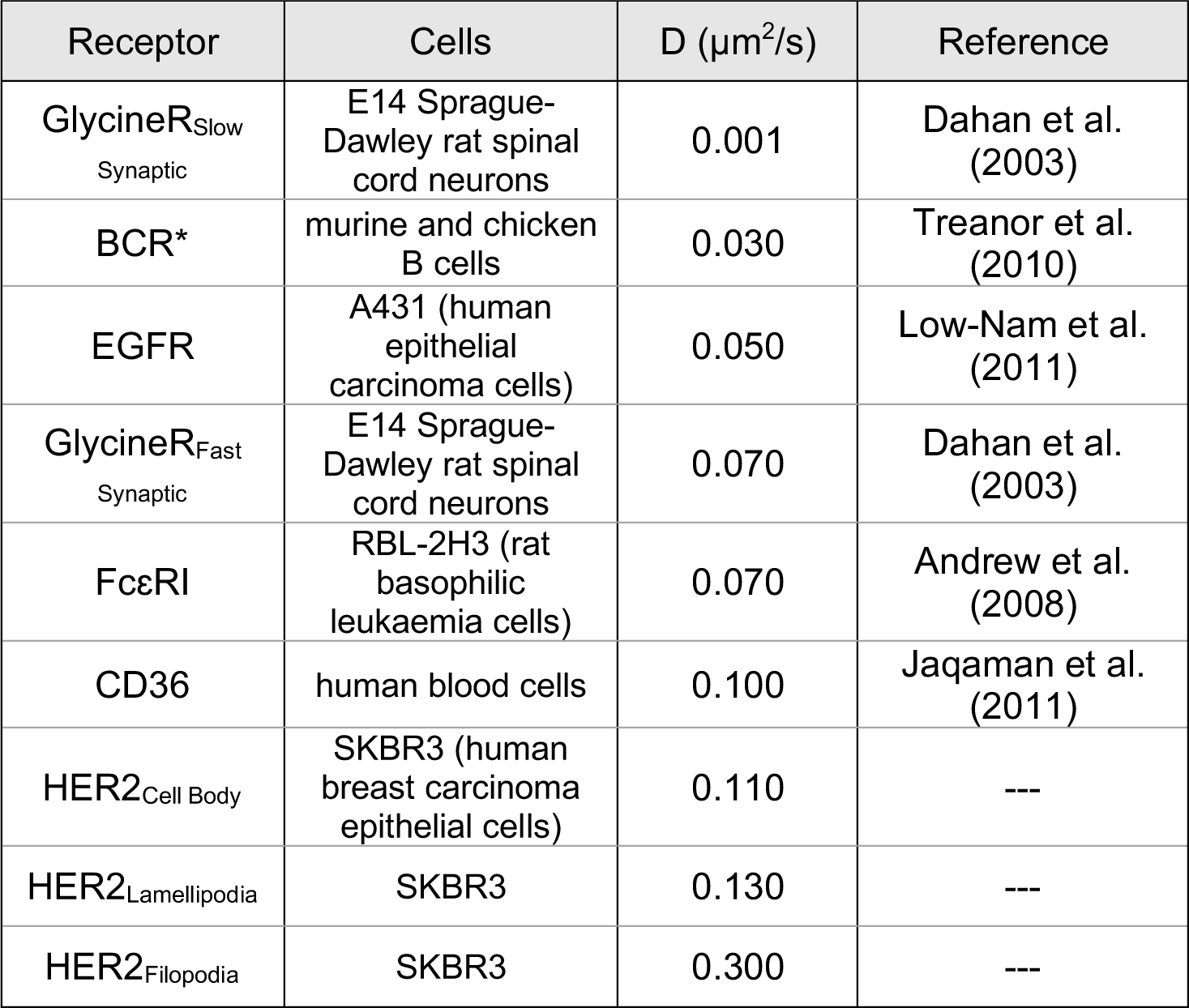
Comparison of diffusion constants of HER2 amongst different cellular regions and other membrane receptor proteins. Comparison of mean HER2 diffusion constant to reported diffusion constants of different membrane proteins. HER2 motion is faster than what has been observed for FcεRI (Andrews et al., 2008), EGFR (Low-Nam et al., 2011), CD36 (Jaqaman et al., 2011), and BCR (Treanor et al., 2010). * Studies cited used single molecule tracking of protein with QDs, with the exception of the BCR study which used single molecule tracking of protein with Cy3.

## Movies

**Movie 1:** DIC videomicroscopy of SKBR3 cells in serum extending protrusions in the form of membrane ruffles, lamellipodia, and filopodia. A motile cell in the upper left corner elaborates a protrusive lamellipodia containing filopodial spikes that is distant by several microns and separated from the cell body. Video acquired at the plane of the coverslip at 1 fps, time denoted as minutes: seconds. Scale bar, 10 µm.

**Movie 2**: Isolated view of lamellipodium and filopodial extension in SKBR3 cell (from **Movie 1**, highlighted in white box of **Figure 1A)**. A lamellipodium extends numerous filopodia and migrates peripherally from the main cell body. Video acquired at the plane of the coverslip at 1 fps and time is in minutes: seconds. Scale bar is 3 µm.

**Movie 3:** Movie of *in vivo* human breast cancer from a biopsy of a metastatic bone lesion as imaged by volume electron microscopy. Individual breast tumor cells have lamellipodia with filopodial protrusions. Within the body of the lamellipodium, mitochondria, small electron lucid vesicles, and rough endoplasmic reticulum was observed. The filopodia extending from the lamellipodium contained actin.

**Movie 4**: Fluorescence of live FITC channel (green) overlaid on a DIC still at a single time point. Localized injection of membrane dye (CellMask™ Plasma Membrane Green) at a subcellular site of SKBR3 cells as a control confirmed localized delivery approximately 5-6 µm from the tip of the pipette immediately following injection. Cells were injected with five 1 sec pulses of dye at t= 6 sec. Live fluorescence video acquired at a plane near the coverslip at 6 fps. Scale bar is 10 µm and time is in minutes: seconds.

**Movie 5**: Protrusion formation occurs at the site of localized, subcellular NRG1 injection of SKBR3 cell. Cells were injected with five 1 sec pulses (over the course of 10 sec) of NRG1 at t= 2 min 50 sec. DIC video acquired at the plane of the coverslip at 1 fps. Scale bar is 10 µm.

**Movie 6**: Continuation of **Movie 5**. Protrusion formation occurs upon second localized, subcellular NRG1 injection of another SKBR3 cell in same preparation as **Movie 5**. Cells were injected with five 1 sec pulses (over the course of 10 sec) of NRG1 at t= 9 min 31 sec. DIC video acquired at a fixed plane at the coverslip at 1 fps. Scale bar is 10 µm.

**Movie 7**: Video of HER2-QD (magenta) overlaid on DIC of SKBR3 cell in serum-containing media. Lamellipodia containing numerous filopodia are observed to undergo continuous ruffling at the leading edge. Though protrusion elaboration is dynamic, HER2-QD motion is distinctly faster than that of protrusions. Note HER2-QDs that appear at the center of cell represent the motion of HER2 on protrusion structures on the ventral surface of the cell at the plane of the substrate. Live video acquired at the plane of the coverslip at 0.5 fps. Scale bar is 10 µm and time is in minutes: seconds.

**Movie 8**: Video of HER2-QD (magenta) overlaid on membrane labeled by WGA-Ax488 in serum-containing media. HER2-QD motion along filopodia extending from the cell body exhibit dynamics that are faster than protrusion elaboration dynamics. Note HER2-QDs that appear at the center of cell represent the motion of HER2 on protrusion structures on the ventral surface of the cell at the plane of the substrate. Live video acquired at the plane of the coverslip at 0.5 fps. Scale bar is 10 µm and time is in minutes: seconds.

**Movie 9**: HER2-QD trajectory on a filopodium displayed as a plot of receptor position over time (*x-y-t*). *x*- and *y*-axes represent position in µm, and *z*-axis represents ascending time in seconds. Different colors on trajectory represent the instantaneous velocity of HER2-QD as specified by the colored scale bar. HER2-QD motion along filopodia was one-dimensionally guided along the length of the protrusion.

**Movie 10**: Video of HER2-QD trajectory on a filopodium (from **Movie 9** and **Figure 4B** Filopodia) overlaid on the HER2-QD channel. Different colors on trajectory represent the instantaneous velocity of HER2-QD as specified by the colored scale bar. Live fluorescence video acquired at the plane of the coverslip at 7 fps, time is in seconds.

**Movie 11**: HER2-QD trajectory on a lamellipodium displayed as a plot of receptor position over time (*x-y-t*). *x*- and *y*-axes represent position in µm, and *z*-axis represents ascending time in seconds. Different colors on trajectory represent the instantaneous velocity of HER2-QD as specified by the colored scale bar. HER2-QD motion along lamellipodia, similar to cell body, was non-directional and consisted of uniform meandering along all axes.

**Movie 12**: Video of HER2-QD trajectory on a lamellipodium (from **Movie 11** and **Figure 4B** Lamellipodia) overlaid on the HER2-QD channel. Different colors on trajectory represent the instantaneous velocity of HER2-QD as specified by the colored scale bar. Live fluorescence video acquired at the plane of the coverslip at 7 fps, time is in seconds.

**Movie 13**: HER2-QD trajectory on cell body displayed as a plot of receptor position over time (*x-y-t*). *x*- and *y*-axes represent position in µm, and *z*-axis represents ascending time in seconds. Different colors on trajectory represent the instantaneous velocity of HER2-QD as specified by the colored scale bar. HER2-QD motion along cell body, similar to lamellipodia, was non-directional and consisted of uniform meandering along all axes.

**Movie 14**: Video of HER2-QD trajectory on cell body (from **Movie 13** and **Figure 4B** Cell Body) overlaid on the HER2-QD channel. Different colors on trajectory represent the instantaneous velocity of HER2-QD as specified by the colored scale bar. Live fluorescence video acquired at the plane of the coverslip at 7 fps, time is in seconds.

**Movie 15**: Latrunculin B (LatB) treatment induced substantial retraction of filopodial protrusions and actin-altered protrusions in SKBR3. Cells were treated with LatB at t= 6 min 36 sec. DIC video acquired at a fixed plane at the coverslip at 1 fps. Scale bar is 10 µm and time is minutes: seconds.

**Movie 16**: Removal of LatB from cells in **Movie 15** and resuspension in media without LatB reversed the depolymerization effects. Cells are observed to slowly re-elaborate protrusions, though protrusions do not regain their fully integrity within the length of acquisition. LatB was removed and cells were resuspended in fresh media (without LatB) prior to the start of video acquisition. DIC video acquired at a fixed plane at the coverslip at 1 fps. Scale bar is 10 µm and time is in minutes: seconds.

**Movie 17**: 3D rendering of actin along the side of a control (-LatB) cell imaged using 3D super-resolution (**Figure 5B**, *left*). Actin-rich filopodial protrusions occur in abundance extending from the cell body, and actin-rich lamellipodial protrusions in the form of membrane ruffles highly decorate the edge of the cell as observed in SEM (see **Figure 2A**). Grid is 2 µm in *x, y*, and *z*.

**Movie 18**: 3D rendering of actin along the side of a +LatB cell imaged using 3D super-resolution (**Figure 5B**, *right*). Cells treated with LatB displayed a substantial alteration in linear actin structure in filopodia and reduced actin-rich filopodial and lamellipodial protrusions. Elaborations present on the membrane appear to be highly retracted ruffles, and filopodial protrusions were absent from the cell surface. Grid is 2 µm in *x, y*, and *z*.

**Movie 19**: HER2-QD trajectory on another filopodia displayed as a plot of receptor position over time (*x-y-t*). *x*- and *y*-axes represent position in µm, and *z*-axis represents ascending time in seconds. Different colors on trajectory represent the instantaneous velocity of HER2-QD as specified by the colored scale bar. HER2-QD motion displayed rapid back-and-forth motion along the length of the filopodium.

**Movie 20**: Video of HER2-QD trajectory on filopodia (from **Movie 19** and **Figure 5C**, -LatB) overlaid on the HER2-QD channel. Different colors on trajectory represent the instantaneous velocity of HER2-QD as specified by the colored scale bar. Live fluorescence video acquired at the plane of the coverslip at 7 fps.

**Movie 21**: HER2-QD trajectory on an actin-altered protrusion in Latrunculin B treated cell displayed as a plot of receptor position over time (*x-y-t*). *x*- and *y*-axes represent position in µm, and *z*-axis represents ascending time in seconds. Different colors on trajectory represent the instantaneous velocity of HER2-QD as specified by the colored scale bar. HER2-QD motion on protrusion of +LatB cell was comparatively less mobile than HER2-QD on filopodia.

**Movie 22**: Video of HER2-QD trajectory on actin-altered protrusion in Latrunculin B treated cell (from **Movie 21** and **Figure 5C**, +LatB) overlaid on the HER2-QD channel. Different colors on trajectory represent the instantaneous velocity of HER2-QD as specified by the colored scale bar. Live fluorescence video acquired at the plane of the coverslip at 7 fps.

## References

1. Pröls, F. Sagar, and M. Scaal, Signaling filopodia in vertebrate embryonic development. Cellular and Molecular Life Sciences, 2016. 73(5): p. 961–974.

2. Gallo, G., Mechanisms underlying the initiation and dynamics of neuronal filopodia: from neurite formation to synaptogenesis. Int Rev Cell Mol Biol, 2013. 301: p. 95–156.

3. Mattila, P.K. and P. Lappalainen, Filopodia: molecular architecture and cellular functions. Nat Rev Mol Cell Biol, 2008. 9(6): p. 446–54.

4. Bettencourt-Dias, M., et al., Centrosomes and cilia in human disease. Trends Genet, 2011. 27(8): p. 307–15.

5. Hildebrandt, F., T. Benzing, and N. Katsanis, Ciliopathies. N Engl J Med, 2011. 364(16): p. 1533–43.

6. Berbari, N.F., et al., The Primary Cilium as a Complex Signaling Center. Current Biology, 2009. 19(13): p. R526–R535.

7. Machesky, L.M., Lamellipodia and filopodia in metastasis and invasion. FEBS Lett, 2008. 582(14): p. 2102–11.

8. Machesky, L.M. and A. Li, Fascin: Invasive filopodia promoting metastasis. Commun Integr Biol, 2010. 3(3): p. 263–70.

9. Arjonen, A., R. Kaukonen, and J. Ivaska, Filopodia and adhesion in cancer cell motility. Cell Adh Migr, 2011. 5(5): p. 421–30.

10. Jacquemet, G., H. Hamidi, and J. Ivaska, Filopodia in cell adhesion, 3D migration and cancer cell invasion. Current Opinion in Cell Biology, 2015. 36: p. 23–31.

11. Wan, L., K. Pantel, and Y. Kang, Tumor metastasis: moving new biological insights into the clinic. Nat Med, 2013. 19(11): p. 1450–64.

12. Menard, S., et al., HER2 overexpression in various tumor types, focussing on its relationship to the development of invasive breast cancer. Ann Oncol, 2001. 12 Suppl 1: p. S15–9.

13. Rubin, I. and Y. Yarden, The basic biology of HER2. Ann Oncol, 2001. 12 Suppl 1: p. S3–8.

14. Yarden, Y. and M.X. Sliwkowski, Untangling the ErbB signalling network. Nat Rev Mol Cell Biol, 2001. 2(2): p. 127–37.

15. Mizutani, T., et al., Relationship of C-erbB-2 protein expression and gene amplification to invasion and metastasis in human gastric cancer. Cancer, 1993. 72(7): p. 2083–2088.

16. Korkaya, H., et al., HER2 regulates the mammary stem/progenitor cell population driving tumorigenesis and invasion. Oncogene, 2008. 27(47): p. 6120–30.

17. Yan, M., et al., HER2 expression status in diverse cancers: review of results from 37,992 patients. Cancer metastasis reviews, 2015. 34(1): p. 157–164.

18. Hommelgaard, A.M., M. Lerdrup, and B. van Deurs, Association with membrane protrusions makes ErbB2 an internalization-resistant receptor. Mol Biol Cell, 2004. 15(4): p. 1557–67.

19. Jeong, J., et al., HER2 signaling regulates HER2 localization and membrane retention. PLOS ONE, 2017. 12(4): p. e0174849.

20. Jeong, J., et al., PMCA2 regulates HER2 protein kinase localization and signaling and promotes HER2-mediated breast cancer. Proc Natl Acad Sci U S A, 2016. 113(3): p. E282–90.

21. Mellman, I. and Y. Yarden, Endocytosis and Cancer. Cold Spring Harbor Perspectives in Biology, 2013. 5(12): p. a016949.

22. Chung, I., et al., High cell-surface density of HER2 deforms cell membranes. Nature Communications, 2016. 7: p. 12742.

23. Lidke, D.S., et al., Reaching out for signals: filopodia sense EGF and respond by directed retrograde transport of activated receptors. The Journal of Cell Biology, 2005. 170(4): p. 619–626.

24. Tada, H., et al., In vivo real-time tracking of single quantum dots conjugated with monoclonal anti-HER2 antibody in tumors of mice. Cancer Res, 2007. 67(3): p. 1138–44.

25. Low-Nam, S.T., et al., ErbB1 dimerization is promoted by domain co-confinement and stabilized by ligand-binding. Nature structural & molecular biology, 2011. 18(11): p. 1244–1249.

26. Jorgens, D.M., et al., Deep nuclear invaginations are linked to cytoskeletal filaments - integrated bioimaging of epithelial cells in 3D culture. J Cell Sci, 2017. 130(1): p. 177–189.

27. Peters, A., et al., Quantitative superresolution microscopy for cancer biology and medicine, in Super-resolution Imaging in Medicine and Biology, A. Diaspro and M.A. van Zandvoort, Editors. 2016, Taylor & Francis Books, Inc.

28. Jacob, T., et al., Ultrasensitive proteomic quantitation of cellular signaling by digitized nanoparticle-protein counting. Sci Rep, 2016. 6: p. 28163.

29. Szent-Gyorgyi, C., et al., Fluorogen-activating single-chain antibodies for imaging cell surface proteins. Nat Biotechnol, 2008. 26(2): p. 235–40.

30. Wang, Y., et al., Affibody-targeted fluorogen activating protein for in vivo tumor imaging. Chem Commun (Camb), 2017. 53(12): p. 2001–2004.

31. Wang, Y., et al., Fluorogen activating protein-affibody probes: modular, no-wash measurement of epidermal growth factor receptors. Bioconjug Chem, 2015. 26(1): p. 137–44.

32. Vermehren-Schmaedick, A., et al., Heterogeneous Intracellular Trafficking Dynamics of Brain-Derived Neurotrophic Factor Complexes in the Neuronal Soma Revealed by Single Quantum Dot Tracking. PLOS ONE, 2014. 9(4): p. e95113.

33. Fichter, K.M., et al., Kinetics of G-protein–coupled receptor endosomal trafficking pathways revealed by single quantum dots. Proceedings of the National Academy of Sciences, 2010. 107(43): p. 18658–18663.

34. Relich, P.K., et al., Estimation of the diffusion constant from intermittent trajectories with variable position uncertainties. Phys Rev E, 2016. 93: p. 042401.

35. Jacquemet, G., et al., FiloQuant reveals increased filopodia density during breast cancer progression. J Cell Biol, 2017. 216(10): p. 3387–3403.

36. Peckys, D.B., U. Korf, and N. de Jonge, Local variations of HER2 dimerization in breast cancer cells discovered by correlative fluorescence and liquid electron microscopy. Science Advances, 2015. 1(6).

37. Hynes, N.E. and H.A. Lane, ERBB receptors and cancer: the complexity of targeted inhibitors. Nat Rev Cancer, 2005. 5(5): p. 341–54.

38. Adam, L., et al., Heregulin regulates cytoskeletal reorganization and cell migration through the p21-activated kinase-1 via phosphatidylinositol-3 kinase. J Biol Chem, 1998. 273(43): p. 28238–46.

39. Courty, S., et al., Tracking individual kinesin motors in living cells using single quantum-dot imaging. Nano Lett, 2006. 6(7): p. 1491–5.

40. Nan, X., et al., Observation of individual microtubule motor steps in living cells with endocytosed quantum dots. J Phys Chem B, 2005. 109(51): p. 24220–4.

41. Sundara Rajan, S. and T.Q. Vu, Quantum Dots Monitor TrkA Receptor Dynamics in the Interior of Neural PC12 Cells. Nano Letters, 2006. 6(9): p. 2049–2059.

42. Trimble, W.S. and S. Grinstein, Barriers to the free diffusion of proteins and lipids in the plasma membrane. J Cell Biol, 2015. 208(3): p. 259–71.

43. Stephen, L.A., Y. Elmaghloob, and S. Ismail, Maintaining protein composition in cilia. Biol Chem, 2017. 399(1): p. 1–11.

44. Nachury, M.V., E.S. Seeley, and H. Jin, Trafficking to the Ciliary Membrane: How to Get Across the Periciliary Diffusion Barrier? Annual Review of Cell and Developmental Biology, 2010. 26(1): p. 59–87.

45. Treanor, B., et al., The membrane skeleton controls diffusion dynamics and signaling through the B cell receptor. Immunity, 2010. 32(2): p. 187–99.

46. Jaqaman, K., et al., Cytoskeletal control of CD36 diffusion promotes its receptor and signaling function. Cell, 2011. 146(4): p. 593–606.

47. Peng, G.E., S.R. Wilson, and O.D. Weiner, A pharmacological cocktail for arresting actin dynamics in living cells. Mol Biol Cell, 2011. 22(21): p. 3986–94.

48. Spector, I., et al., Latrunculins: novel marine toxins that disrupt microfilament organization in cultured cells. Science, 1983. 219(4584): p. 493.

49. Huang, F.K., et al., Targeted inhibition of fascin function blocks tumour invasion and metastatic colonization. Nat Commun, 2015. 6: p. 7465.

50. Yamaguchi, H. and J. Condeelis, Regulation of the actin cytoskeleton in cancer cell migration and invasion. Biochimica et biophysica acta, 2007. 1773(5): p. 642–652.

51. Mattila, P.K., F.D. Batista, and B. Treanor, Dynamics of the actin cytoskeleton mediates receptor cross talk: An emerging concept in tuning receptor signaling. J Cell Biol, 2016. 212(3): p. 267–80.

